# Modelling the spatial-temporal distributions and associated determining factors of a keystone pelagic fish

**DOI:** 10.1101/2020.04.16.044156

**Authors:** Samantha Andrews, Shawn J. Leroux, Marie-Josée Fortin

## Abstract

Mobile pelagic species habitat is structured around dynamic oceanographic and ecological processes which operate and interact horizontally and vertically throughout the water column and change over time. However, pelagic species movements and distributions are often poorly understood. We use the Maxent species distribution model to assess how changes in the relative importance of modelled oceanographic (e.g., temperature) and climatic variables (e.g., the North Atlantic Oscillation) over 17-years affect the monthly average horizontal and vertical distribution of a keystone pelagic forage species, Atlantic Canadian capelin (*Mallotus villosus*). We show the range and distribution of capelin occurrence probabilities vary across horizontal and vertical axes over time, with binary presence/absence predictions indicating capelin occupy between 0.72% (April) and 3.45% (November) of the total modelled space. Furthermore, our analysis reveals that the importance of modelled oceanographic variables, such as temperature, vary between months (44% permutation importance in August to 2% in May). By capturing the spatial dynamics of capelin over horizontal, vertical, and temporal axes, our analysis builds on work that improves our understanding and predictive modelling ability of pelagic species distributions under current and future conditions for pro-active ecosystem-based management.

## Introduction

Mobile pelagic species have critical functions in marine systems, occupying multiple trophic levels (Sarà and Sarà, 2007; van der Lingen *et al.*, 2010), acting as agents for resource flows (Gounand *et al.*, 2018), and as ecosystem engineers (Breitburg *et al.*, 2010). Despite this, their extensive movements which can cross jurisdictional boundaries means they are typically poorly surveyed. Consequently, the abundance, distribution, and population structures of mobile pelagic species are not well understood.

Species distribution models (SDMs), which combine abiotic (e.g. temperature) and biotic (e.g. prey) variables with species presence data, are commonly used to predict potential species distributions (Guisan *et al.*, 2017). SDMs have been applied to a wide range of marine species including invertebrates (Eger *et al.*, 2017), reef fish (Young and Carr, 2015), and sponges (Knudby *et al.*, 2013). Modelling pelagic species distributions remains challenging, however, because pelagic species habitat is structured around dynamic oceanographic and ecological processes, operating and interacting horizontally and vertically throughout the water column, and changing over time (Angel, 1993). Oceanographic conditions that influence pelagic species distributions include temperature (Rose, 2005), salinity (Vilhjálmsson, 2002), dissolved oxygen content (Bertrand *et al.*, 2011), sea surface height (Zainuddin *et al.*, 2017), and mixed layer depth (Williams *et al.*, 2015). Prey abundance and distribution also plays a pivotal role in pelagic predator distributions (Garrison *et al.*, 2002), with chlorophyll concentration often acting as a proxy for zooplanktivorous prey (Bailey *et al.*, 2012). Furthermore, large-scale climate oscillations such as the North Atlantic Oscillation and Atlantic Multidecadal Oscillation influence regional oceanographic conditions and have been implicated in changing distributions of pelagic species (Gregory *et al.*, 2009).

While distribution models are typically applied to two-dimensional surfaces which lack temporal variability, the dynamic horizontal, and vertical fluidity of the ocean necessitates models – particularly those generated for mobile pelagic species which can be distributed throughout the water column – to account for conditions in four-dimensions – across longitude and latitude, at depth, and over time. Whereas oceanographic data averaged over long temporal windows (e.g. decadal, annual) may provide sufficient information to model the distributions of relatively sedentary species, using such data can lead to loss of information about how pelagic species are responding spatially to variations in conditions than more contemporaneous resolutions (e.g. monthly or daily) (Mannocci *et al.*, 2017).

Distribution models that consider changes through time are increasing in number (e.g. Brodie *et al.*, 2018). However, although the availability in-situ and remotely sensed oceanographic data and ocean models that provide data at multiple depths is increasing (Kavanaugh *et al.*, 2016), distribution models that incorporate conditions at depth are rare (Duffy and Chown, 2017, but see Hobday and Hartmann, 2006). Such oceanographic information is not truly three-dimensional, but rather offers snapshots at specific depths creating a “2.5-dimensional” environment (Duffy and Chown, 2017). Nevertheless, incorporating and explicitly modelling both depth and time into SDMs may improve our ability to predict current and future mobile pelagic species distributions and explore biological responses to global change.

Capelin (*Mallotus villosus*) is a migratory zooplanktivorous pelagic fish species with a circumpolar distribution in the Pacific, Arctic, and North Atlantic Oceans. Four genetically distinct populations exist – two in the Pacific, one around West Greenland, and one in the Northeast Atlantic around Atlantic Canada (Præbel *et al.*, 2008). Within Atlantic Canadian waters, capelin, whose abundance and distribution is closely tied to environmental conditions, is regarded as a keystone species (Rose, 2005; Davoren *et al.*, 2006). During the 1990s, the North Atlantic experienced a substantial change in ocean climate (Greene *et al.*, 2008). Within Atlantic Canadian waters, alongside numerous groundfish species capelin declined in abundance, shifted their distributions, began maturing at younger ages and smaller sizes, and spawned later in the year (Carscadden and Nakashima, 1997). Capelin have not recovered to their former state, and these changes have been implicated in the lack of recovery of Canada’s Atlantic cod (*Gadus morhua*), which previously had significant economic importance in the region (Mullowney and Rose, 2014). Despite the ecological and cultural importance of capelin in Atlantic Canada, their year-round distributions and spatial responses to changing conditions throughout the region is not fully understood.

In this study, we address the following research objectives: (1) quantify variation in the average monthly distribution of capelin across longitude, latitude, and depth to capture their horizontal and vertical movements, and (2) assess changes in the relative importance of modelled oceanographic and climatic variables to estimated capelin distributional changes. We construct monthly “2.5-dimensional” distribution models with Maxent (Phillips *et al.*, 2006), predict the average monthly estimated probability of occurrence across the 2.5-dimensional geographic space, and derive the relative importance of modelled variables using Maxent’s permutation importance metrics. Results of our analysis will broadly contribute to the application of distributions models for mobile pelagic species, and more specifically to the understanding of capelin in Atlantic Canada, a forage species with substantial ecological and cultural importance in the region.

## Methods

### Study region

Our study region includes parts of the Atlantic Canadian ocean domain - the ocean and benthos lying within and adjacent to the Canadian territorial sea (0 – 12 nautical miles from the low-water line along the coast) and Exclusive Economic Zone (EEZ) (12 – 24 nautical miles from the low-water line along the coast), broadly extending from 40° latitude to 70° latitude. To avoid imposing unnatural breaks in capelin distributions, we include France’s EEZ surrounding Saint Pierre and Miquelon, disputed territories with the USA, and areas adjacent to Canada’s EEZ such as the Flemish Cap and Georges Bank and Basin as part of the study area (Figure 1).

**Figure 1:**
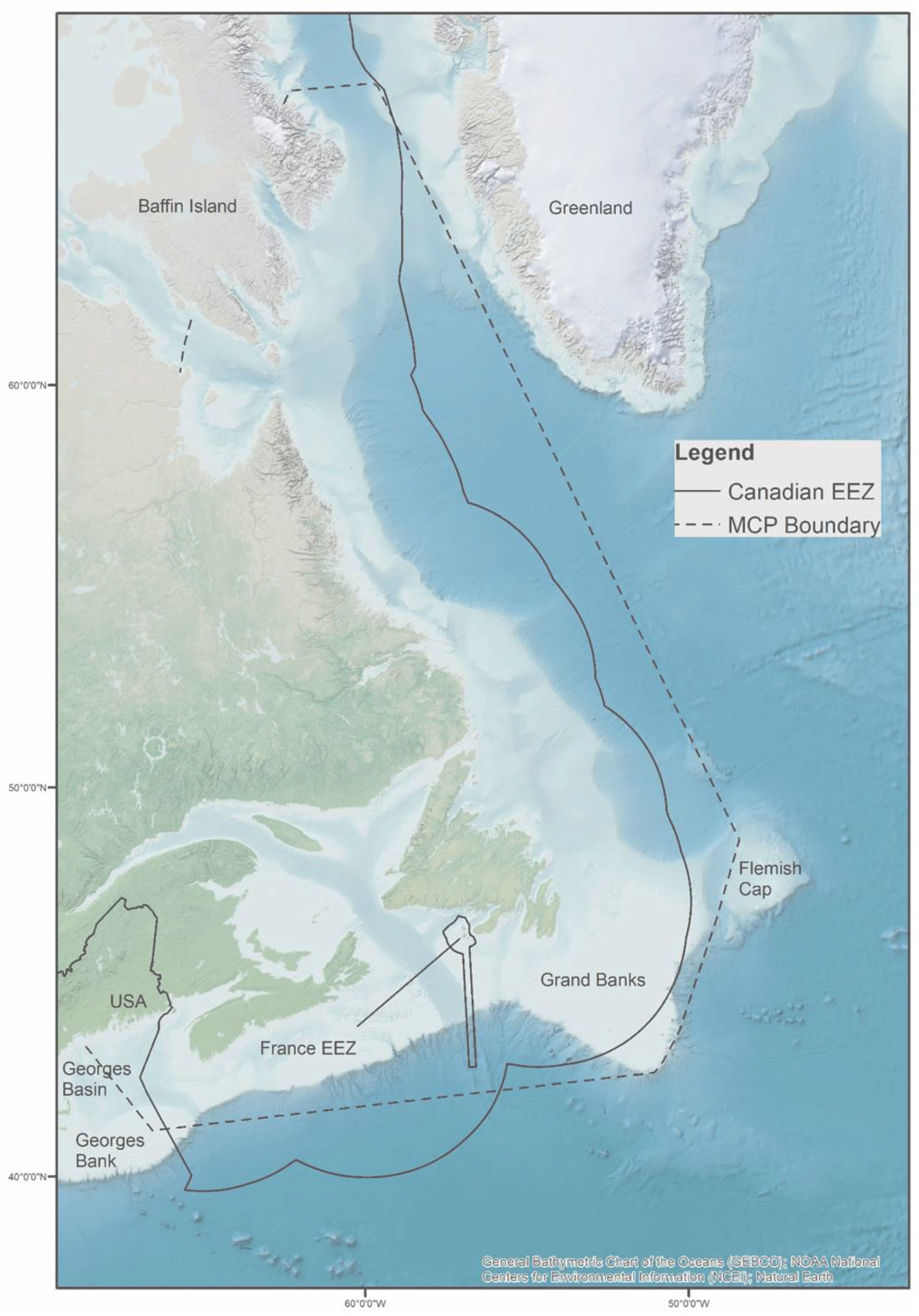
The study region includes waters inside the Canadian EEZ (purple line), the French EEZ contained within the Canadian EEZ, and adjacent areas of the Flemish Cap, Grand Banks, Georges Bank, and Georges Basin. Analysis was restricted to the Minimum Convex Polygon (MCP) boundaries, which was derived from capelin presence data.

### Data – Environmental variables

We obtained oceanographic data (chlorophyll α concentration, density mixed layer depth, dissolved oxygen, salinity, sea surface height, and temperature), as well as bottom depth, from the numerical modelling products GLORYS V4.1 (E.U. Copernicus Marine Service Information, 2018a) and BIOMER V3.2 (E.U. Copernicus Marine Service Information, 2018b). Both products offer global gridded monthly average measurements at a horizontal resolution of 0.25° across 75 depth levels (0.5 meters to 5,902 meters at varying intervals) for the years 1998 to 2015 inclusive.

We also obtained monthly averages for two climate oscillations; the Atlantic Multidecadal Oscillation (AMO) index (ESRL, 2019) calculated from detrended spatially averaged sea surface temperature anomalies between 87.5° - 87.5°N, 2.5° - 357.5°E and the North Atlantic Oscillation (NAO) index (NCAR, 2019) calculated from the leading empirical orthogonal function sea level pressure anomalies between 20°-80°N, 90°W-40°E. In both cases, index values vary temporally but not spatially across the study region (i.e. characterises broad temporal variability in ocean conditions within and between months).

### Data – Capelin presences

We obtained georeferenced capelin presence data from the Ocean Biogeographic Information System (OBIS) database (OBIS, 2018). We subset the dataset to the years 1998 to 2014 to match the oceanographic data availability (we excluded 2015 as there were only ten presences recorded in the OBIS dataset for the entire year) and only included data points with sampling depth information. To reduce over-representation of conditions that may arise from over-sampling of cells (Elith *et al.*, 2006), we reduced the number of presences to one per spatial-temporal grid cell (see below for details on the grid). For example, if a cell had twelve presence points during March 1999, we reduced the number of presences in that cell to one.

For individual species and size classes, sampling gears differ in their ability to catch individuals, influencing detectability of presences and potentially biasing SDMs (Knudby *et al.*, 2013). We gleaned sampling gear information from the metadata submitted to OBIS from data contributors to include as a categorical predictor. Based on the metadata, we identified nine different, albeit broad, gear type categories (Supplementary File 1).

### Spatial-temporal grid

To account for the 3-dimensional nature of the ocean in the SDMs, we created a series of grids representing each depth included in the model, based on the oceanographic data layers. Because our distribution model (i.e., Maxent, see below) assumes an equal-area surface, we projected all layers into an Albers Equal Area (25 km^2^) grid for modelling. The grid follows this same projection. The grid was duplicated to create one grid per month-year period, thus giving each grid cell a unique *x*, *y*, *z*, and *t* location. Throughout the remainder of this text, grids which include only a spatial element will be referred to as the ‘spatial grid’ while the grid that includes the month-year period will be referred to as the ‘spatial-temporal grid’.

### Modelling Process

We used Maxent (Phillips *et al.*, 2006 Supplementary File 2) to model the average spatial distribution of capelin presence probabilities on a month-by-month basis (e.g. one model for March including March data from all years between 1998 and 2014 for which there is observation data). We carried out all analysis in R version 3.5.3 (R Core Team, 2019), with the *Raster* (Hijmans, 2019), *Dismo* (Hijmans *et al.*, 2017), *EcoSpat* (Broennimann *et al.*, 2018), and *enmSdm* (Smith, 2019) packages.

### Background Point Generation

Maxent uses background points to characterise the study area. To reduce model over-fitting and to accurately reflect conditions across a species’ range, background points should be constrained to the geographical range of known occurrence (Elith *et al.*, 2011; Merow *et al.*, 2013). We derived the capelin’s horizontal geographic range using a 100% minimum convex polygon (MCP) (Syfert *et al.*, 2014) (Figure 1). To include depth with the MCP, we also excluded depth layers falling below the maximum depth of capelin presences (maximum depth layer 1,045.85 meters). We generated 10,000 background points for each monthly model (Supplementary File 3). Only one background point per spatial-temporal grid cell was permitted to prevent over-representation of oceanographic conditions (Elith *et al.*, 2006).

To account for between-month variability (e.g. March 1999 vs March 2000) in oceanographic conditions, we weighted the number of background points to the number of presences recorded in each month-year period. For example, if 22% of the unique observations by cell occurred in 2003, 49% in 2004, 7% in 2005, and 21% in 2006, 22% of the background points for that model came from 2003, 49% from 2004, 7% in 2005, and 21% in 2006. To ensure we sufficiently captured conditions across the capelin’s range, we also weighted points by the number of spatial grid cells represented in each Northwest Atlantic Fisheries Organization (NAFO) division and the area of the Hudson Strait represented in the MCP (NAFO, 2019). For example, we subdivided the 22% of the background points that came from 2003 so that 8.8% of the cells came from NAFO division 3K, 7.4 from 3L, 4% from 3Ps, etc. (Supplementary File 4). This double-weighting of background points also acts to reduce the influence of sampling bias, which may impact the performance of the models (Elith et al., 2011).

### Data extraction for presence and background points

We extracted ten oceanographic, six climatic, and one static (bathymetry) environmental predictor variables (Table 1) to the presence and background points. We included both surface values and values at the depth layer closest to capelin presence/background points as predictors (Supplementary File 5) except for sea surface height, density mixed layer depth, seafloor depth, the AMO, and the NAO as these are single layer values only. To account for the potential lagged influence of AMO and NAO on distributions, we added three metrics for the AMO and NAO predictors: the value of the oscillations during the sampling month, the value during the previous sampling month, and the mean value from the previous winter (calculated from December to February values).

**Table 1:**
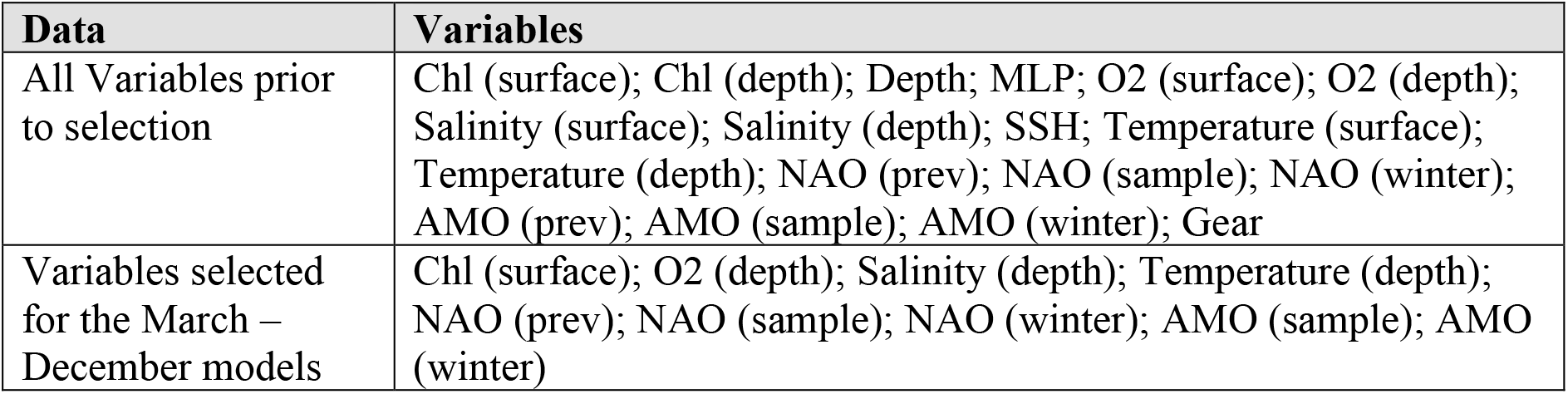
Variables used before and after a-priori variable selection based on Spearman Correlation and VIF for the March – December models. Variable abbreviations: depth – value of the variable at presence depth; surface – value of the variable at the sea surface; Chl – Chlorophyll concentration (mmol.m ^−^3); Depth – Sea floor depth; MLP - Mixed layer thickness (meters); O2 – Dissolved Oxygen (mmol.m ^−^3); Salinity – Salinity (PSU); SSH- Sea Surface Height (meters) Temperature – Temperature (kelvin); NAO – North Atlantic Oscillation; AMO – Atlantic Multidecadal Oscillation; sample – value during the presence month; prev – value from the month prior to presence; winter – value during the winter previous to presence; Gear – Gear Type (Table SF1 in supplementary materials).

| Data | Variables |
| --- | --- |
| All Variables prior to selection | Chl (surface); Chl (depth); Depth; MLP; O<sub>2</sub> (surface); O<sub>2</sub> (depth); Salinity (surface); Salinity (depth); SSH; Temperature (surface); Temperature (depth); NAO (prev); NAO (sample); NAO (winter); AMO (prev); AMO (sample); AMO (winter); Gear |
| Variables selected for the March – December models | Chl (surface); O<sub>2</sub> (depth); Salinity (depth); Temperature (depth); NAO (prev); NAO (sample); NAO (winter); AMO (sample); AMO (winter) |

### Variable selection

To minimise the impact of collinearity, we selected variables for each monthly model a-priori using Spearman correlation coefficient and variance inflation factor (VIF) (Dormann *et al.*, 2013) (Supplementary File 6). Although Maxent performs well even with small sample sizes (Pearson *et al.*, 2007), to avoid erroneous estimation of predictor variance, we employed a minimum events per variable (EPV) rule in which we would only model months that had a minimum of five samples per predictor (Yalcin and Leroux, 2018).

### Maxent modelling

We executed individual monthly models using Maxent version 3.4.1. To allow entropy to be reached, we set the maximum number of iterations to 5000 for each model. We also set regularization (β) multiplier values for each monthly model to reduce overfitting and complexity. To determine the regularization values, we constructed a series of models for each month with regularization values ranging from 0.25 to 4, at 0.25 step intervals. Optimal regularization multiplier values were determined using Akaike’s Information Criterion adjusted for small sample size (AICc) (Warren and Seifert, 2011). Final models for each month used the regularization value that resulted in the lowest AICc.

For each monthly model we used temporally-split cross-validation to train and test the model (Radosavljevic and Anderson, 2014), in which data from one year was used as the training data, and tested against all other years, and then repeated until we used all years for training. We further assessed models performance using several metrics - testing and training area under the curve (AUC) scores (Phillips *et al.*, 2006), true skills statistic (TSS) (Allouche *et al.*, 2006), and the continuous Boyce index (CBI) (Boyce *et al.*, 2002) (Supplementary File 7). We calculated each metric separately for each temporal fold which we then averaged to produce mean predictive performances for each monthly model. Finally, we created permutation plots for each monthly model to measure mean variable importance of each of the environmental variables used in the model.

### Spatial predictions

For each month, we created average probability maps based on Maxent’s cloglog transformation, which provides an estimated probability of presence (Phillips *et al.*, 2017). In order to predict estimated probability of occurrence across the depth layers, we concatenated the depth layers of each oceanographic variables into a single stacked variable layer for each temporal fold (Bentlage *et al.*, 2013). For example, for March 1999, we had one layer for chlorophyll α concentrations representing all depth layers, one layer for dissolved oxygen, one for salinity, and one for temperature. We used these stacked layers to produce probability maps for each temporal fold, which we then averaged to create monthly average probability maps for each depth-layer (Supplementary File 8).

To explore general patterns in the probability of capelin presence, we also converted the probability maps into binary presence/absence maps using a maximum Kappa derived threshold. This threshold is used in a wide range of distribution modelling studies (e.g. (Davidson *et al.*, 2017; Scherrer *et al.*, 2018), demonstrates resiliency to prevalence which cannot be accurately determined with presence-only data, and shows good agreement with the AUC (Nenzén and Araújo, 2011).

## Results

We obtained a total of 11,517 presence points that matched the required spatial and temporal extent from OBIS. Of these, we removed 5,157 presences that lacked depth information, were duplicate entries or appeared in the same spatial-temporal cell. A total of 6,360 presence points remained. Presence point data availability was highly variable by month, with June having the highest number of presences (*n* = 1,263) and February the lowest number of presences (*n* = 10) throughout the study period (1998-2014) (Supplementary File 9).

Of the eighteen variables selected a-priori, multicollinearity analysis indicated only six, five, and nine variables could be included in the final models for January, February, and March to December respectively. However, following the EVP rule, we did not run a February model as there were insufficient presence points. We also chose to not to run a January model due to the smaller number of sampling years compared to the others (four years vs eleven+ years for March – December). Selected variables for the March to December models are presented in Table 1.

### Maxent model results

Each of the monthly models on average performed very well (metrics Train AUC, Test AUC, TSS, and CBI; Supplementary File 7). Overall, mean performance of the models was highest in December and lowest in October.

The average (across temporal folds) relative contribution of the variables to each monthly model also varied (Figure 2). Temperature contributed the most to predicting capelin distribution in the June (41%), July (27%), August (44%), September (41%), October (46%), and November (31%) models, and the least to the March (11%), April (3%), and May (2%) models. Salinity had the strongest influence in the March model (38%); however, during no month did it contribute the least to a model fit. Chlorophyll α had the strongest influence on predicted distribution in the December model (41%) and was least important in the May (5%), June (9%), August (1%), and September (3%) models. Dissolved oxygen was the most important predictor of capelin distribution in the May model (51%), and the least in the July (10%), October (8%), and December (12%) models. The climate oscillations (NAO and AMO) played little role in any of the models, nor did they vary substantially in their importance from one month to the next (range 0.06 for NAO value from the previous month in June, to 16% for the NAO value from the previous winter in April).

**Figure 2:**
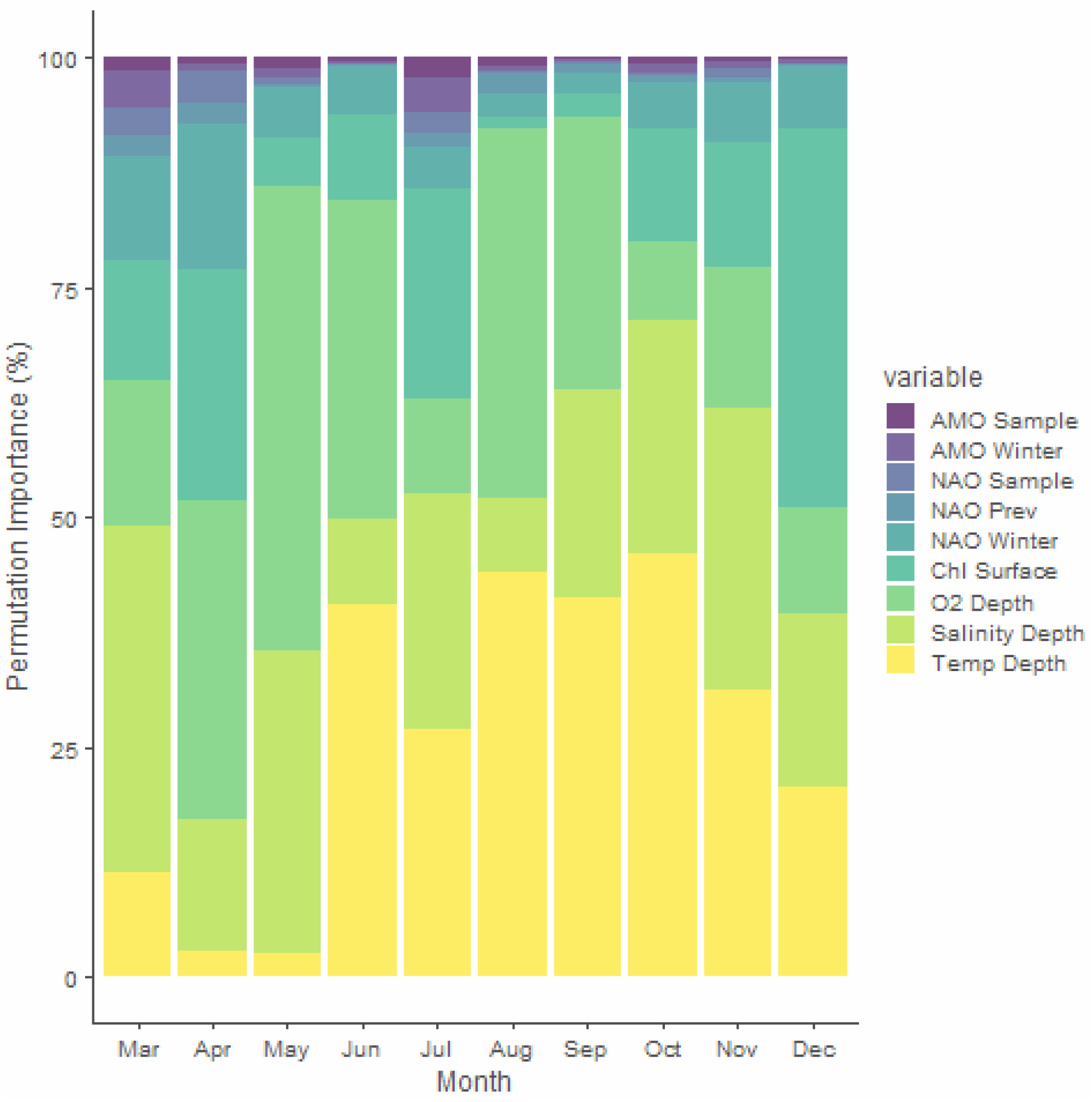
Average permutation importance of environmental variables used in the Maxent models. Values represent averages (across temporal folds) for each month. See Table 1 for a definition and description of the variables

### Monthly probability maps

We produced a total of 460 average (across temporal folds) prediction maps based on Maxent’s cloglog transformation, depicting the probability of capelin occurrence from ~0.5 meters depth to ~1,045 meters depth between March and December inclusive (Figure 3).

**Figure 3:**
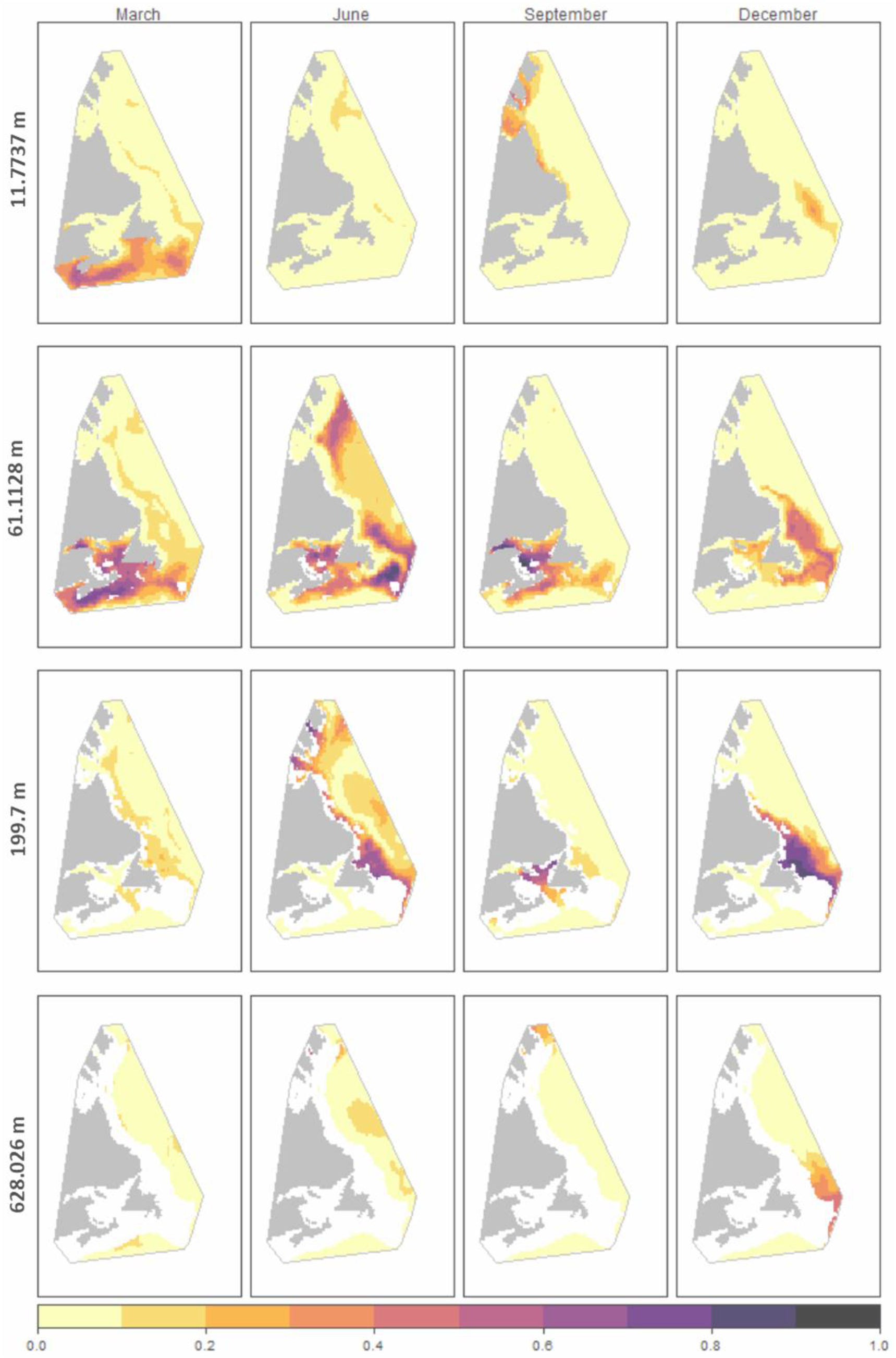
Sample of predicted probability of occurrence presence maps generated by Maxent for March, June, September, and December over four different depth layers. White indicates the seafloor and grey indicates landmass. Lighter colours indicate lower probabilities and darker colours higher.

Alongside latitudinal and longitudinal distributional changes, the average vertical distribution of probabilities of occurrence also varied by month (Figure 4). On average, depths of ~244 meters and shallower saw the highest probability scores, except for October, November, and December which saw higher probabilities at greater depths. Across all depths, the spread of probability values varies from month to month though some general patterns emerge when binning the probabilities into ten equally sized groups (Table 2). Most probability values fall below 0.3 (65.3% in May to 94.07% in September). Between 0.3 and 1, values fall steadily to under 1% of the total number of spatial grid cells. There are two exceptions to this pattern. In both August and October, there is a small increase in probabilities falling between 0.7 and 0.79, though values continue to decline thereafter.

**Figure 1:**
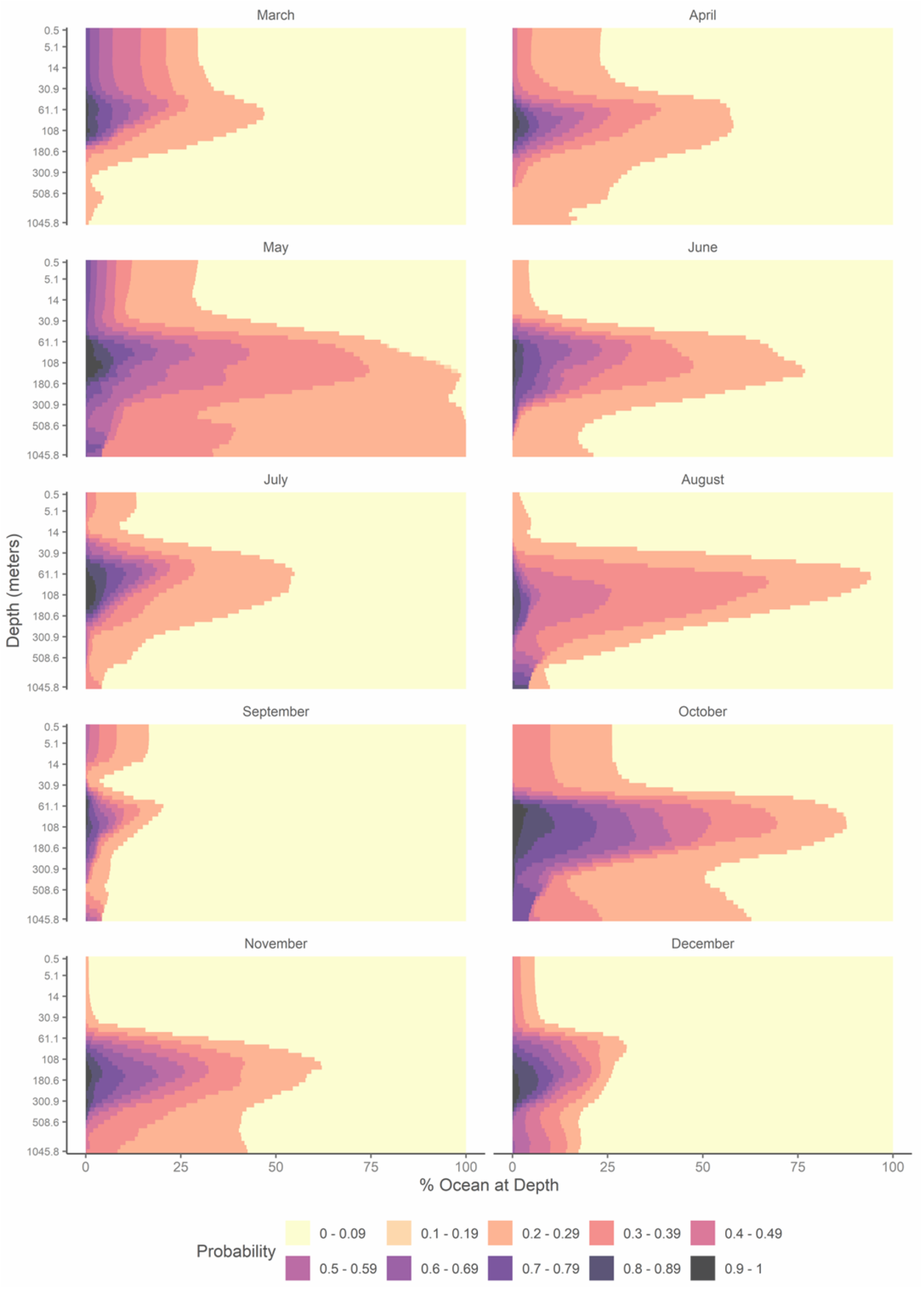
Vertical distribution of Maxent model predicted probability of capelin occurrence from March to December relative to the amount of ocean available at each depth. Note that there is a greater area of ocean at higher depths than lower depths due to increasing intrusion of the sea floor, and that depth layers are not evenly spaced (see Supplementary File 5). Lighter colours indicate lower probability and darker colours higher.

When converting the probability maps into binary predicted presence/absence, we see differences in the horizontal (*x*, *y*) and vertical (*z*) distribution of capelin predicted presences across the study area (Figures 5 and 6). From March to June, predicted presence cells shifted eastwards, extending out from the Gulf of St Lawrence and around Nova Scotia to encompass eastern Newfoundland and the Grand Banks (western trailing cell centroid in March −67.75° to −64.5° in June, eastern leading cell centroid in March −55.25° to −46.25° in June), with some cells appearing further north. The vertical distribution of predicted presence cells also expanded during this period, with cells occupying depth layers from ~0.5 meters to ~147 meters in March to ~42 meters to ~656 meters in June. From July to September, predicted presence cells largely shrunk back to the Gulf of St Lawrence (western trailing cell centroid in July −69.25° to −69° in September, eastern leading cell centroid in July −52.5° to −57.5° in September), before expanding again in October to encompass the eastern side of Nova Scotia and Newfoundland. During August, predicted presence cells occupied the greatest number of depth layers (~27 meters to ~1046 meters). In November predicted cells show a shift eastwards (western trailing cell centroid in October −62.0° to −59.5° in November, eastern leading cell centroid in October −48° to −46.75° in November) and northwards (southern trailing cell centroid in October 43° to 44.75° in November, northern leading cell centroid in October - to −57.5° in September) along Labrador before contracting to eastern Newfoundland in December. The number of depth layers with predicted presences also reduced during this period (~69 meters to ~509 meters in November, and ~69 meters to ~458 meters in December). Following these distribution patterns, the percentage spatial grid cells predicted to contain capelin in all months remains relatively small – from 0.72% of the total spatial grid in April up to a maximum of 3.45% in August (Figure 5).

**Table 2:** Percentage of spatial grid cells falling into Maxent predicted probability of capelin occurrence bins for each of the modelled months. For example, in March 71.88% of cells had a predicted probability of occurrence between 0 and 0.09.

|  | Mar | Apr | May | Jun | Jul | Aug | Sep | Oct | Nov | Dec |
| --- | --- | --- | --- | --- | --- | --- | --- | --- | --- | --- |
| **0 - 0.09** | 71.88 | 64.49 | 36.52 | 67.70 | 71.70 | 60.29 | 88.21 | 47.87 | 75.13 | 84.48 |
| **0.1 - 0.19** | 0 | 0.01 | 0.18 | 0.01 | 0 | 0 | 0 | 0.03 | 0 | 0 |
| **0.2 - 0.29** | 11.96 | 22.03 | 28.61 | 16.87 | 17.18 | 17.91 | 5.86 | 22.58 | 10.56 | 4.72 |
| **0.3 - 0.39** | 4.90 | 6.54 | 16.89 | 5.94 | 4.32 | 14.43 | 2.47 | 11.70 | 5.48 | 3.45 |
| **0.4 - 0.49** | 4.46 | 2.95 | 6.64 | 3.20 | 1.82 | 5.15 | 1.36 | 4.27 | 2.85 | 2.29 |
| **0.5 - 0.59** | 2.62 | 1.69 | 4.97 | 2.34 | 1.42 | 0.72 | 0.79 | 3.34 | 2.26 | 1.83 |
| **0.6 - 0.69** | 2.08 | 1.05 | 3.08 | 1.94 | 1.23 | 0.46 | 0.55 | 3.78 | 1.83 | 1.18 |
| **0.7 - 0.79** | 1.34 | 0.58 | 1.73 | 1.32 | 1.02 | 0.56 | 0.47 | 4.02 | 1.18 | 0.95 |
| **0.8 - 0.89** | 0.62 | 0.45 | 0.81 | 0.49 | 0.85 | 0.47 | 0.21 | 1.73 | 0.52 | 0.90 |
| **>0.9** | 0.14 | 0.22 | 0.58 | 0.18 | 0.45 | 0.03 | 0.08 | 0.69 | 0.18 | 0.21 |

**Figure 5:**
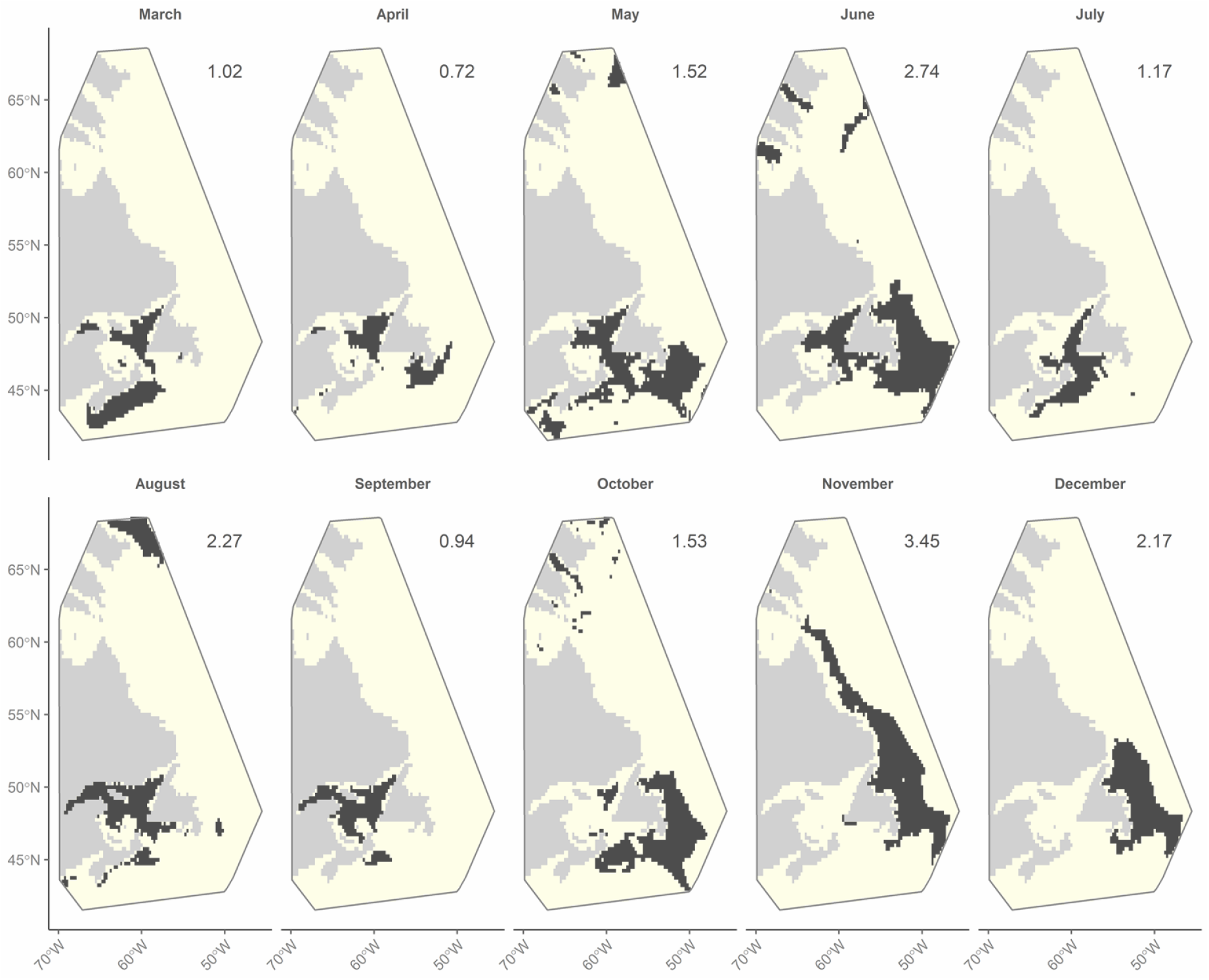
2-Dimensional Overview of the horizontal (x, y) average location of predicted capelin presence (black) and absence (yellow) for each modelled month, based on maximum kappa thresholds applied to Maxent model outputs. Areas of occurrence indicates the location of a presence cell, regardless of depth. The value on the top right indicated the percentage of cells in the spatial grid with predicted presence.

**Figure 6:**
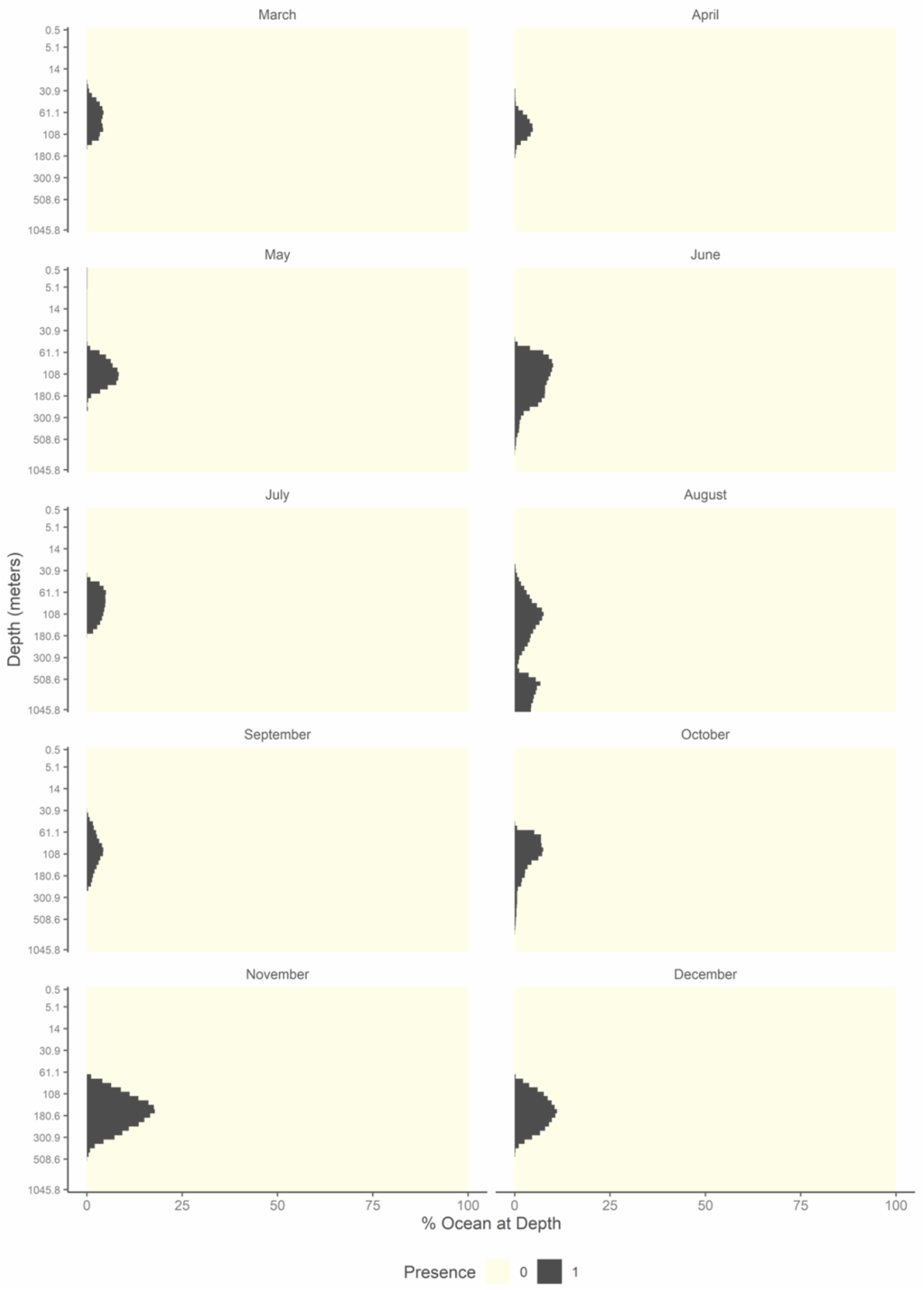
The vertical average distribution of predicted capelin presence (black) and absence (yellow) relative to the amount of ocean available at each depth layer for each modelled month, based on maximum kappa thresholds. Sea surface is represented at the top of the y-axis and deepest depth at the bottom. Note that there is a greater area of ocean at higher depths than lower depths due to increasing intrusion of the sea floor, and that depth layers are not evenly spaced (see Supplementary File 5).

## Discussion

Marine-based SDMs are typically applied to ‘flat’ surfaces, lacking temporal and vertical information. However, for marine pelagic species which can exhibit extensive movements horizontally (*x, y*) and vertically (*z*) throughout the water column, models should explicitly consider space in four dimensions – across depth and time as well as longitude and latitude. By coupling long time-series (17 years) of species presence data with “2.5-dimensional” oceanographic models, we quantified the spatial-temporal dynamics of the estimated probability of capelin occurrence throughout the water column across Atlantic Canada, and relative importance of modelled variables on a month-by-month basis.

Our analysis indicates the range and distribution of capelin occurrence probabilities vary month by month, and over depth, with the highest probability scores for March – September occurring in shallower depths than for October - December. When binarizing probability values into presence/absence, we found capelin occupy between 0.72% (April) and 3.45% (November) of the total modelled space. We also observed depth changes with, for example, presences primarily located further west in shallower waters in March (~ 26 to ~133 meters) than in December (~78 to ~371 meters). This pattern is consistent with observations of capelin moving eastwards and into deeper waters towards the end of the year (Carscadden and Nakashima, 1997; Davoren *et al.*, 2006). While alternative thresholds may adopt a lower value for binarizing predictions into presence/absence (Nenzén and Araújo, 2011), we expect the general distribution patterns to remain.

By producing monthly models, we were able to assess the relative importance of modelled variables for monthly capelin distribution. Our analysis revealed that chlorophyll α concentrations, dissolved oxygen, salinity, and temperature varied in importance as predictors from one month to the next, with each becoming the most important variable in at least one of the months modelled. Notably temperature contributed the most to the June – November models. Temperature is widely considered a key determinate of marine fish distribution, including for capelin (Rose, 2005), though has also been found to play a small role in interannual variations in the distribution of some species (Thorson *et al.*, 2017). Along similar lines, we found temperature contributed the least to the March - May and December models. Alongside temperature, oxygen places substantial metabolic constraints on marine species distributions (Deutsch *et al.*, 2015), a pattern which emerged in the April and May models in which dissolved oxygen became the most important predictor of capelin occurrence.

The changes in relative variable importance revealed in our models may indicate capelin’s realized niche changes relative to variations in conditions experienced in each month and/or with phenology. Indeed, the distribution and frequency of modelled variable values vary across the modelled space. For example, predictably water temperatures are on average warmer in August than in March, and chlorophyll α concentrations higher in May than in August (Supplementary Files 3 and 10). Furthermore, the physiology of any given species determines which suite of conditions are suitable for survival. While species have upper and lower limits to physiological tolerances, species may experience conditions that vary within their tolerance range over time (Stuart-Smith *et al.*, 2017). By capturing changes in variable importance for pelagic species, models have the potential more closely reflect temporal dynamics in species realized niche, improving our understanding and predictive ability of changing distributions under current and future conditions. Additionally, given prey location plays a role in predator distributions (Garrison et al., 2002), incorporating the spatial-temporal dynamics of capelin distributions may serve to better inform models for capelin predator distributions and multispecies assessments in ecosystem-based fishery management.

Distribution models require consideration of the relevance of the spatial and temporal scales at which species and oceanographic processes operate on (Mannocci *et al.*, 2017). The oceanographic models chosen to model distributions dictate the resolution of our models (monthly averaged 25° grid cell). While the oceanographic models allowed us to encompass a large time-period of capelin data, the resolution may prove too coarse particularly during biologically important periods, such as during the spawning period when capelin physiology and behaviour changes (Rose, 2005; Davoren *et al.*, 2006). Furthermore, the models presented in this study provide monthly average distributions, potentially masking interannual variation in conditions that determine distributions (Boyce *et al.*, 2002). Nevertheless, the continued development of finer spatial and temporal scale models may not only allow for more refined distribution models, but their predictive output can allow for increased efficiency in the implementation of management measures (Lewison *et al.*, 2015; Dunn *et al.*, 2016), including nowcasting and forecasting of species distributions for pro-active management.

Our study adds to the small albeit growing body of distribution models which embrace a “2.5 dimension” approach, and models that attempt to capture distribution changes on timescales more relevant to the dynamic nature of pelagic species. With survey effort spatially and temporally patchily distributed throughout our modelled space, our predicted occurrences offer new insights into the potential distribution of capelin - a forage species with substantial ecological importance in Atlantic Canada, and whose life-histories are sensitive to changing environmental conditions. By capturing species spatial-temporal dynamics over horizontal and vertical axes, and changes in the relative importance of modelled variables, such studies can enhance our understanding and predictive modelling ability of pelagic species distributions under current and future conditions for pro-active ecosystem-based conservation and fisheries management.

## Supplementary Material

The following supplementary material is available:

Supplementary File 1: Gear type categories identified in the OBIS metadata.
Supplementary File 2: Brief background on Maxent
Supplementary File 2: Brief background on Maxent.
Supplementary File 3: Selecting the number of background points to use in each monthly model.
Supplementary File 4: NAFO division with percentage weighting for background point creation.
Supplementary File 5: Depth layers used to model and predict probability of capelin occurrence.
Supplementary File 6: Variable Selection: Spearman Correlation and VIF Tables.
Supplementary File 7: Model performance metrics.
Supplementary File 8: Concatenating depth layers into single raster files for prediction.
Supplementary File 9: Year-month distribution of capelin presence points.
Supplementary File 10: Variation in Environmental Correlates.

## Acknowledgements

This research is sponsored by the Natural Sciences and Engineering Research Council of Canada’s Canadian Healthy Oceans Network and its Partners: Department of Fisheries and Oceans Canada and the Institut Nordique de Recherche en Environnement et en Santé au Travail (representing the Port of Sept-Îles and City of Sept-Îles). The authors wish to thank Fran Mowbray (DFO-Newfoundland), the Ecosystem Ecology Lab (Memorial University), Paul Snelgrove (Memorial University), Mariano Kohen-Alonso (DFO-Newfoundland), and Tom Therriault (DFO-Pacific) for their feedback throughout the process. We also thank a number of people who have helped with the underlying code used to create the analysis in this study: David Bazin (Copernicus Marine Environment Monitoring Service); Michael D. Sumner (Australian Antarctic Division); Rémi Daigle (DFO-Maritimes); Simon Beaton (Independent), and the wider Stack Exchange Community, with particular thanks to: TinglTanglBob (Stack Overflow); Nadizan (Stack Overflow); Spacedman (GIS Stack Exchange).

## Data Availability Statement

All datasets used in this study were derived from sources in the public domain and are cited in the text. In order of citation these datasets are:

GLORYS V4.1: E.U. Copernicus Marine Service Information. 2018a. Global Ocean Reanalysis Products (GLOBAL-REANALYSIS-PHY-001-025) Version 4.1. Available from https://marine.copernicus.eu/services-portfolio/access-to-products/.
BIOMER V3.2: E.U. Copernicus Marine Service Information. 2018b. Global Biogeochemical Non assimilative Hindcast Product (GLOBAL_REANALYSIS_BIO_001_018) Version 3.2. Available from https://marine.copernicus.eu/services-portfolio/access-to-products/.
The Atlantic Multidecadal Oscillation (AMO) index: ESRL. 2019. Atlantic Multidecadal Oscillation (AMO) Index (Kaplan SST V2). https://www.esrl.noaa.gov/psd/data/timeseries/AMO/ (Accessed 2 January 2019).
North Atlantic Oscillation (NAO) index: NCAR. 2019. Hurrell North Atlantic Oscillation (NAO) Index (PC-Based). https://climatedataguide.ucar.edu/climate-data/hurrell-north-atlantic-oscillation-nao-index-pc-based (Accessed 2 January 2019).
Capelin presence: OBIS. 2018. Distribution records of Mallotus villosus (Lamarck, 1798). Available: Ocean Biogeographic Information System. Intergovernmental Oceanographic Commission of UNESCO. www.iobis.org (Accessed 31 October 2018).

