## Supplementary File 1 for "Modelling the spatial-temporal distributions and associated determining factors of a keystone pelagic fish"

### Supplementary File 1: Gear type categories identified in the OBIS metadata.

Table SF1: Gear type information is provided by data contributors when submitting their datasets to the Ocean Biogeographic Information System (OBIS) database. There is no reporting standard for gear type and as such, details provided by contributors varied between datasets. We removed gear type from due to insufficient gear information and concerns that some reporting may have led to incorrect gear type classification.

| **Gear Type** | **Frequency** |
|---|---|
| Bottom trawl "Alfredo-3" | 3 |
| Bottom trawl "Campelen-14" | 51 |
| Bottom trawl "Campelen-1800" | 7689 |
| Bottom trawl "Campelen-21" | 56 |
| Bottom trawl "Cosmos-2600" | 62 |
| Bottom trawl "Western IIA" | 1220 |
| Bottom trawl (unknown) | 652 |
| Vertical plankton tow | 19 |
| Not specified | 547 |

NOT PEER REVIEWED
