## Supplementary File 2 for "Modelling the spatial-temporal distributions and associated determining factors of a keystone pelagic fish"

### Supplementary File 2: Brief background on Maxent.

Maxent (Phillips *et al.*, 2006) is a widely-used presence-background model based on the maximum entropy principle – that subject to prior knowledge, the probability distribution which best describes the data is that with maximum entropy (i.e. the least informative distribution). By comparing known presences with the background environment, Maxent's output indicates the extent to which the model fits presence data more or less than it would if the presences had a uniform distribution. Maxent performs well compared to the arguably more robust presence-absence based models such as generalized linear models (GLMs) and generalized additive models (GAMs), including with small sample sizes when regularization (β) multiplier values are tuned to the model (Elith *et al.*, 2006; Pearson *et al.*, 2007).

NOT PEER REVIEWED
