## Supplementary File 3 for "Modelling the spatial-temporal distributions and associated determining factors of a keystone pelagic fish"

### Supplementary File 3: Selecting the number of background points to use in each monthly model.

Although as a general rule Maxent performance does not substantially improve with more than 10,000 background points, some studies involving large study areas such as ours have found using a much larger number of background points that more fully capture the variation in conditions across a species range results in better performing models (Guevara *et al.*, 2018). To determine the optimal number of background points whilst still allowing for efficient computational processing time, we compared the distribution of each oceanographic variable collected from 10,000, 20,000, 50,000, 100,000, and 190,000 randomly generated points from three different time periods – February 1999, June 2014, and October 2007 (Figures SF3.1, SF3.2, and SF3.3). As there was little difference between the distributions, in the interests of efficiency we opted to generate 10,000 background points per model.

NOT PEER REVIEWED

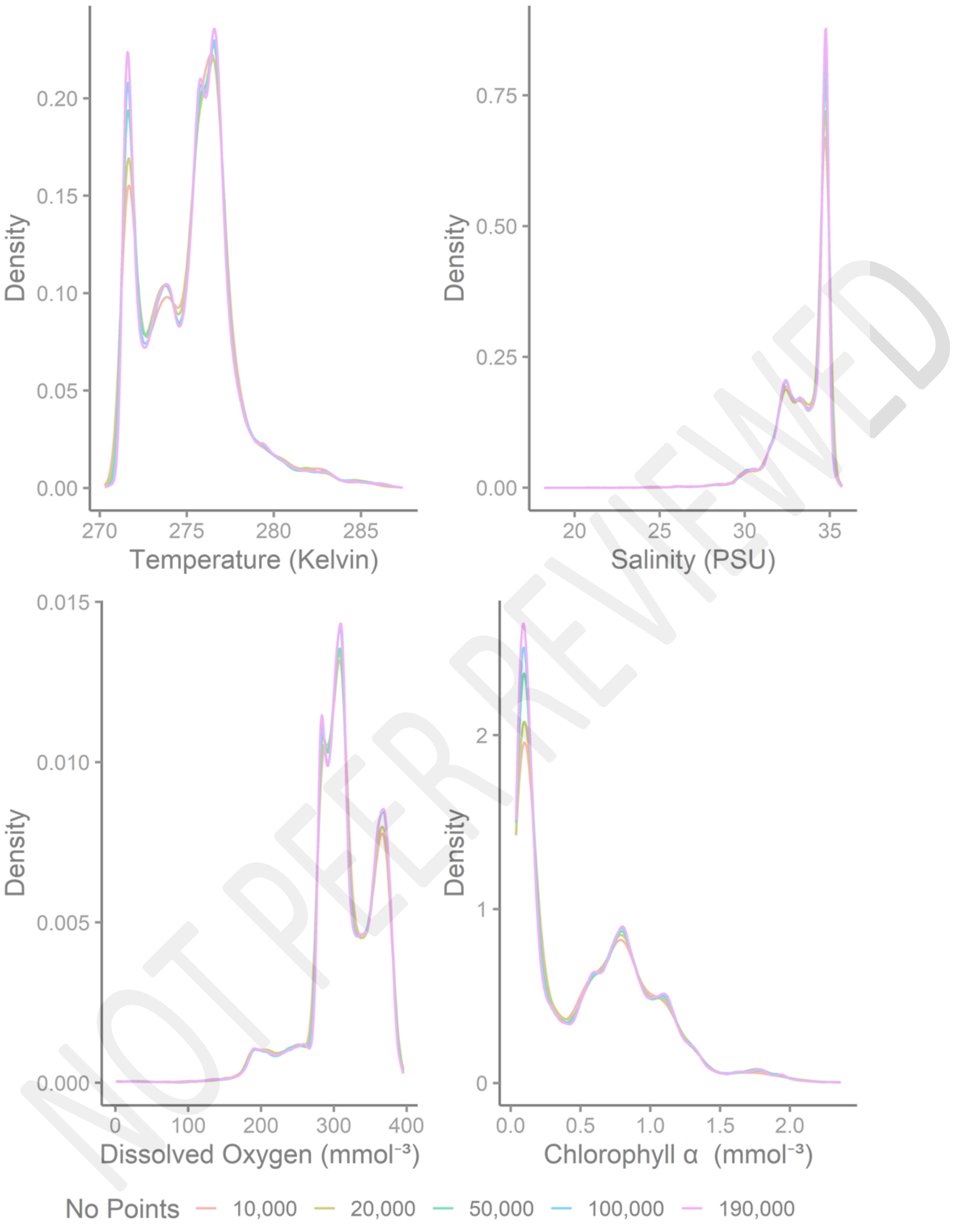


Figure SF3.1: Distribution of each oceanographic variable collected from 10,000, 20,000, 50,000, 100,000, and 190,000 randomly generated points during February 1992.

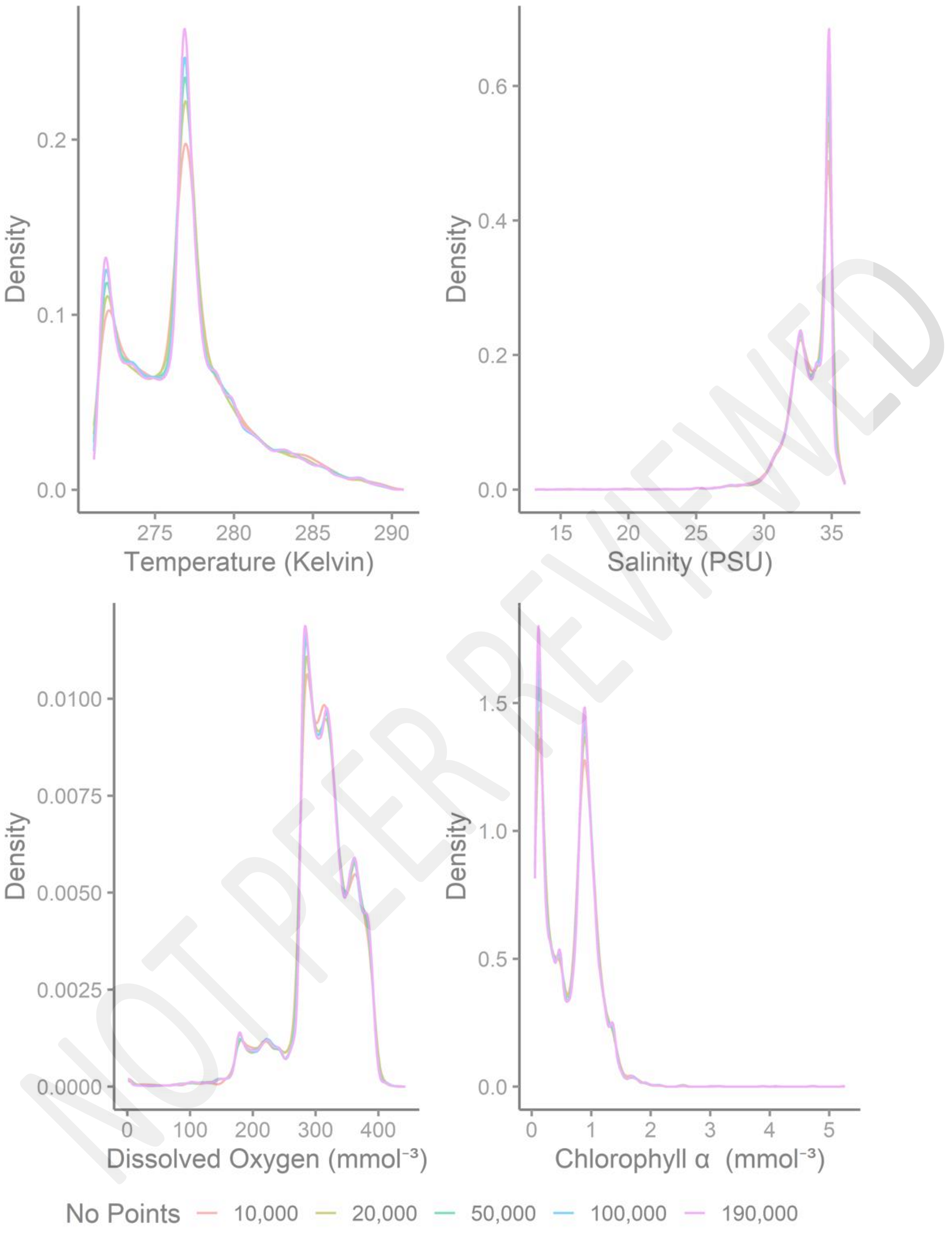


Figure SF3.2: Distribution of each oceanographic variable collected from 10,000, 20,000, 50,000, 100,000, and 190,000 randomly generated points during June 2004.

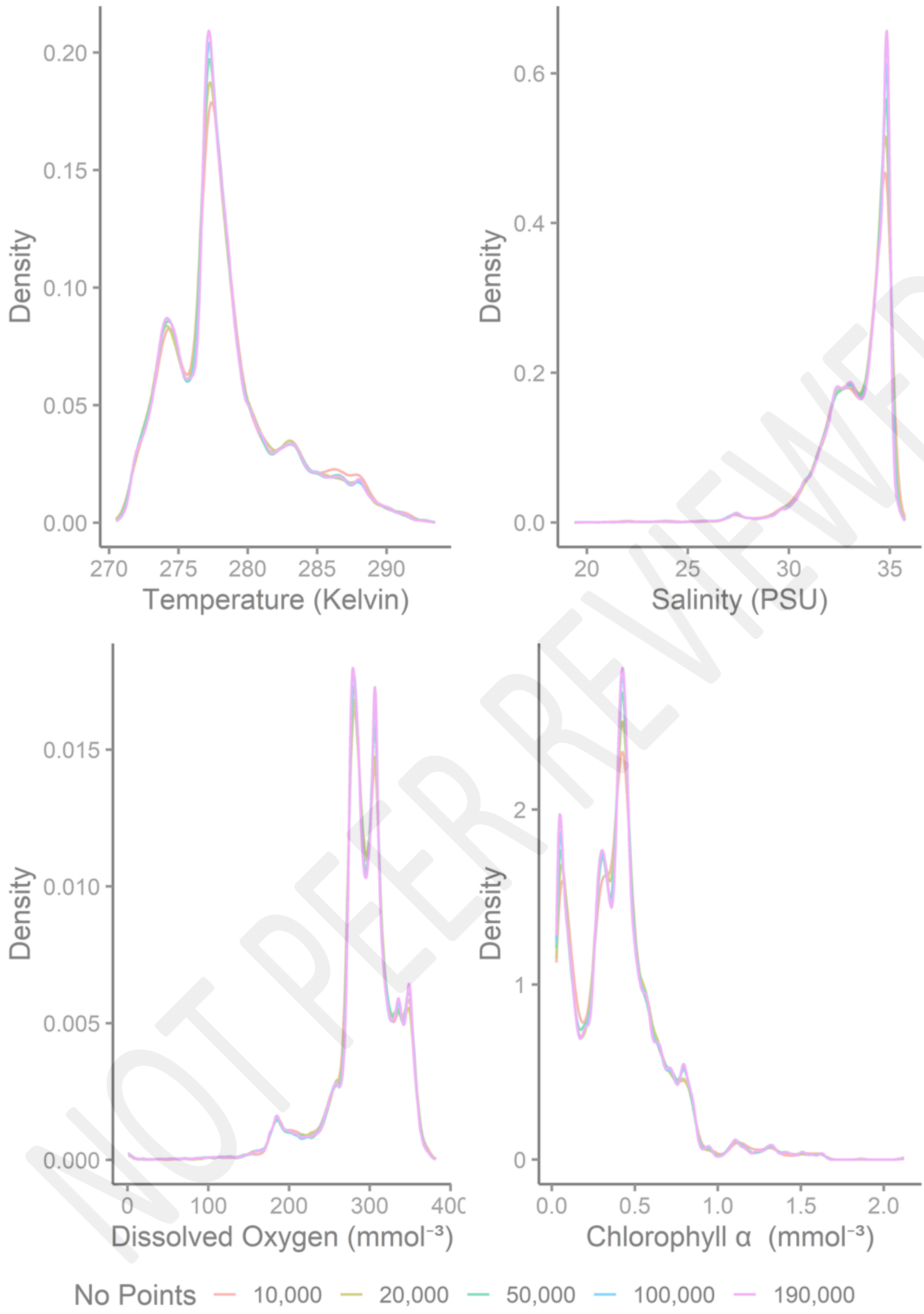

NOT PEER REVIEWED

Figure SF3.3: Distribution of each oceanographic variable collected from 10,000, 20,000, 50,000, 100,000, and 190,000 randomly generated points during October 2007.
