## Supplementary File 4 for "Modelling the spatial-temporal distributions and associated determining factors of a keystone pelagic fish"

### Supplementary File 4: NAFO division with percentage weighting for background point creation.

To ensure that we sufficiently captured conditions across the capelin's range, we also weighted points by the number of spatial grid cells represented in each Northwest Atlantic Fisheries Organization (NAFO) division and the area of the Hudson Strait represented in the MCP (Figure SF4).

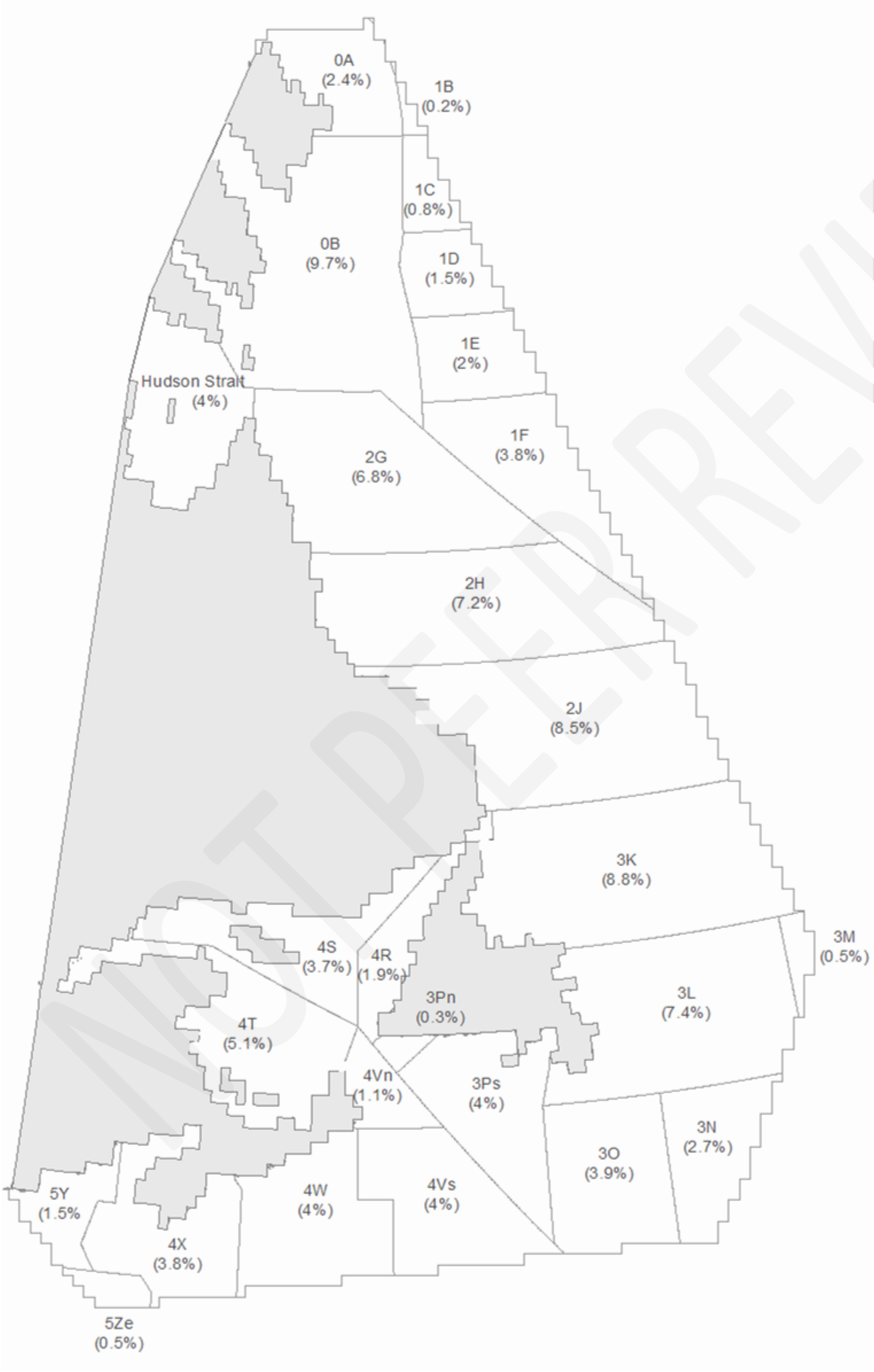


Figure SF4: NAFO divisions and percentage of spatial grid cells within each division. Map created in ESRI ArcMap 10.5.

NOT PEER REVIEWED
