## Supplementary File 5 for "Modelling the spatial-temporal distributions and associated determining factors of a keystone pelagic fish"

### Supplementary File 5: Depth layers used to model and predict probability of capelin occurrence.

The GLORYS and BIOMER ocean models are not truly 3-dimensional. Instead they represent the global ocean as a series of depth layers (Table SF5), with each layer providing information as to the conditions at that particular layer. Because conditions are not available at exactly the same depth as capelin samples came from, we used the depth layer closest to the sampling depth. For example, is a sample came from a depth of 127 meters, we used conditions from the depth layer representing conditions at 133.08 meters as this layer is closer than the depth layer representing 120 meters.

Table SF5: The Minimum Convex Polygon (MCP) encompasses 46 depth layers derived from the GLORYS (E.U. Copernicus Marine Service Information, 2018a) and BIOMER (E.U. Copernicus Marine Service Information, 2018b) ocean models. Depth values have been rounded to two decimal points.

| Layer No | Depth (meters) | Layer No | Depth (meters) |
|---|---|---|---|
| 1 | 0.51 | 24 | 97.04 |
| 2 | 1.56 | 25 | 108.03 |
| 3 | 2.67 | 26 | 120.00 |
| 4 | 3.86 | 28 | 133.08 |
| 5 | 5.14 | 29 | 163.17 |
| 6 | 6.54 | 30 | 180.55 |
| 7 | 8.09 | 31 | 199.79 |
| 8 | 9.82 | 32 | 221.14 |
| 9 | 11.77 | 33 | 244.89 |
| 10 | 13.99 | 34 | 271.36 |
| 11 | 16.53 | 35 | 300.89 |
| 12 | 19.43 | 36 | 333.86 |
| 13 | 22.76 | 37 | 370.69 |
| 14 | 26.56 | 38 | 411.79 |
| 15 | 30.87 | 39 | 457.63 |
| 16 | 35.74 | 40 | 508.64 |
| 17 | 41.18 | 41 | 565.29 |
| 18 | 47.21 | 42 | 628.03 |
| 19 | 53.85 | 43 | 697.26 |
| 20 | 61.11 | 44 | 773.39 |
| 21 | 69.02 | 45 | 856.68 |
| 22 | 77.61 | 46 | 947.45 |
| 23 | 86.93 | 47 | 1045.85 |

NOT PEER REVIEWED

NOT PEER REVIEWED
