## Supplementary File 6 for "Modelling the spatial-temporal distributions and associated determining factors of a keystone pelagic fish"

### Supplementary File 6: Variable Selection: Spearman Correlation and VIF.

To minimise the impact of collinearity, we selected variables for each monthly model a-priori using Spearman correlation coefficient and variance inflation factor (VIF) (Dormann *et al.*, 2013). We removed variables with Spearman's correlation coefficients above 0.7 (Table SF6.1) and VIF values above 5, (Table SF6.2) and gave variable retention preference to oceanographic variables closest to sampling depth as, due to proximity, they likely have a more direct impact on species occurrence than more distal values.

Table SF6.1: Spearman correlation coefficients for all variables indicating highly (> 0.7 / < -0.7 - red) and moderately (> 0.5 / < -0.5 - blue) correlated variables.

| **Variable1** | **Variable2** | **Jan** | **Feb** | **Mar** | **Apr** | **May** | **Jun** | **Jul** | **Aug** | **Sep** | **Oct** | **Nov** | **Dec** |
|---|---|---|---|---|---|---|---|---|---|---|---|---|---|
| AMO (sampling month) | AMO (previous winter) | -0.97 | 1 | - | - | - | - | - | - | - | - | - | - |
| AMO (sampling month) | AMO (previous month) | 1 | 0.85 | 0.64 | 0.69 | 0.82 | 0.87 | 0.89 | 0.94 | 0.92 | 0.73 | 0.94 | 0.71 |
| AMO (sampling month) | NAO (sampling month) | -0.78 | -0.85 | 0.53 | - | -0.65 | - | - | - | - | - | - | - |
| AMO (sampling month) | NAO (previous winter) | -0.54 | 1 | -0.64 | - | - | - | - | - | - | - | - | - |
| AMO (sampling month) | NAO (previous month) | 0.73 | 1 | - | - | -0.59 | -0.62 | - | 0.57 | - | -0.54 | - | - |
| AMO (previous winter) | AMO (previous month) | -0.97 | 0.85 | - | - | - | - | - | - | - | - | - | - |
| AMO (previous winter) | NAO (sampling month) | 0.86 | -0.85 | - | - | - | - | - | - | - | - | - | - |

NOT PEER REVIEWED

| | | | | | | | | | | | | | |
|---|---|---|---|---|---|---|---|---|---|---|---|---|---|
| AMO (previous winter) | NAO (previous winter) | - | -1 | - | - | - | - | - | - | - | - | - | - |
| AMO (previous winter) | NAO (previous month) | -0.55 | 1 | - | - | - | - | - | - | - | 0.54 | - | - |
| AMO (previous month) | NAO (sampling month) | -0.78 | -1 | - | - | 0.5 | - | - | - | - | - | - | - |
| AMO (previous month) | NAO (previous winter) | -0.54 | -0.85 | -0.7 | - | -0.57 | - | - | - | - | - | - | - |
| AMO (previous month) | NAO (previous month) | 0.73 | 0.85 | - | - | - | -0.56 | - | - | - | - | - | - |
| NAO (sampling month) | NAO (previous winter) | 0.55 | 0.85 | - | - | - | - | - | - | - | - | - | - |
| NAO (sampling month) | NAO (previous month) | - | -0.85 | - | - | - | - | 0.55 | - | - | - | - | - |
| NAO (previous winter) | NAO (previous month) | -0.86 | -1 | - | -0.53 | - | - | - | - | 0.51 | - | - | - |
| Chl (surface) | Chl (depth) | 0.51 | 0.5 | - | - | - | - | - | - | - | - | - | - |
| Chl (surface) | Temp (surface) | - | - | - | - | -0.55 | -0.76 | -0.69 | -0.66 | -0.72 | -0.77 | - | - |
| Chl (surface) | Temp (depth) | - | - | - | - | - | -0.57 | - | - | - | -0.54 | - | - |
| Chl (surface) | O2 (surface) | - | - | - | - | 0.52 | 0.8 | 0.74 | 0.68 | 0.76 | 0.82 | 0.52 | - |
| Chl (surface) | O2 (depth) | - | - | - | - | - | - | - | - | - | 0.51 | - | - |
| Chl (surface) | Salinity (surface) | -0.52 | - | - | - | - | - | - | - | - | - | - | - |
| Chl (depth) | O2 (depth) | 0.66 | 0.69 | 0.72 | 0.76 | 0.77 | 0.79 | 0.72 | 0.67 | 0.64 | 0.67 | 0.64 | 0.6 |
| Chl (depth) | Salinity (depth) | -0.72 | -0.7 | -0.67 | -0.63 | -0.56 | -0.62 | -0.59 | -0.55 | -0.56 | -0.64 | -0.72 | 0.73 |
| Mixed Layer Depth | Sea Surface Height | -0.82 | -0.73 | -0.71 | -0.72 | -0.73 | -0.74 | -0.7 | -0.65 | -0.71 | -0.77 | -0.77 | -0.78 |

NOT PEER REVIEWED

| | | | | | | | | | | | | | |
|---|---|---|---|---|---|---|---|---|---|---|---|---|---|
| Mixed Layer Depth | O2 (surface) | -0.65 | -0.62 | -0.61 | -0.57 | - | - | - | - | - | - | - | -0.5 |
| Mixed Layer Depth | Salinity (surface) | 0.86 | 0.76 | -0.78 | 0.81 | 0.81 | 0.84 | 0.75 | 0.73 | 0.83 | 0.86 | 0.86 | 0.85 |
| Mixed Layer Depth | Salinity (depth) | 0.65 | 0.57 | 0.56 | 0.6 | 0.57 | 0.59 | 0.53 | 0.5 | 0.58 | 0.62 | 0.64 | 0.65 |
| Mixed Layer Depth | Bottom depth | -0.65 | -0.59 | -0.59 | -0.6 | -0.59 | -0.61 | -0.6 | -059 | -0.66 | -0.67 | -0.67 | -0.66 |
| Sea Surface Height | O2 (surface) | 0.70 | 0.79 | 0.74 | 0.71 | - | - | - | - | - | - | - | - |
| Sea Surface Height | Salinity (surface) | -0.95 | -0.96 | -0.94 | -0.93 | -0.92 | -0.89 | -0.88 | -0.88 | -0.85 | -0.9 | -0.91 | -0.93 |
| Sea Surface Height | Salinity (depth) | -0.76 | -0.79 | -0.75 | -0.77 | -0.74 | -0.7 | -0.67 | -0.63 | -0.6 | -0.66 | -0.7 | -0.74 |
| Sea Surface Height | Bottom depth | 0.74 | 0.78 | 0.75 | 0.76 | 0.75 | 0.75 | 0.71 | 0.70 | 0.68 | 0.71 | 0.73 | 0.75 |
| Temp (surface) | Temp (depth) | 0.89 | 0.85 | 0.84 | 0.86 | 0.82 | 0.73 | 0.63 | 0.54 | 0.63 | 0.71 | 0.84 | 0.89 |
| Temp (surface) | O2 (surface) | -0.80 | -0.74 | -0.68 | -0.68 | -0.84 | -0.95 | -0.96 | -0.97 | -0.97 | -0.97 | -0.94 | -0.86 |
| Temp (surface) | O2 (depth) | -0.56 | -0.53 | -0.51 | -0.5 | - | -0.52 | -0.54 | -0.55 | -0.62 | -0.66 | -0.64 | -0.59 |
| Temp (surface) | Salinity (depth) | - | 0.5 | 0.5 | - | - | - | - | - | - | - | - | - |
| Temp (surface) | Bottom depth | -0.53 | -0.59 | -0.56 | - | - | - | - | - | - | - | - | - |
| Temp (depth) | O2 (surface) | -0.76 | -0.67 | -0.64 | -0.69 | -0.81 | -0.75 | -0.65 | 0.57 | -0.65 | -0.72 | -0.83 | -0.84 |
| Temp (depth) | O2 (depth) | -0.73 | -0.79 | -0.78 | -0.73 | -0.61 | -0.5 | -0.52 | -0.51 | -0.55 | -0.53 | -0.57 | -0.66 |
| Temp (depth) | Salinity (depth) | 0.58 | 0.64 | 0.67 | 0.6 | - | - | - | - | - | - | - | - |
| Temp (depth) | Bottom depth | -0.52 | -0.58 | -0.56 | -0.54 | - | - | - | - | - | - | - | - |
| O2 (surface) | O2 (depth) | 0.63 | 0.6 | 0.59 | 0.59 | 0.62 | 0.57 | 0.59 | 0.6 | 0.65 | 0.7 | 0.7 | 0.68 |
| O2 (surface) | Salinity (surface) | -0.68 | -0.78 | -0.79 | -0.75 | - | - | - | - | - | - | - | - |
| O2 (surface) | Salinity (depth) | -0.66 | -0.74 | -0.73 | -0.69 | - | - | - | - | - | - | - | - |
| O2 (surface) | Bottom depth | 0.66 | 0.77 | 0.77 | 0.73 | 0.51 | - | - | - | - | - | - | 0.5 |
| O2 (depth) | Salinity (depth) | -0.75 | -0.77 | -0.78 | -0.77 | -0.7 | -0.62 | - | - | - | - | 0.6 | -0.68 |
| O2 (depth) | Bottom depth | 0.53 | 0.57 | 0.57 | 0.56 | - | - | - | - | - | - | - | - |

NOT PEER REVIEWED

| | | | | | | | | | | | | | |
|---|---|---|---|---|---|---|---|---|---|---|---|---|---|
| Salinity (surface) | Salinity (depth) | 0.78 | 0.8 | 0.78 | 0.81 | 0.78 | 0.76 | 0.73 | 0.72 | 0.73 | 0.74 | 0.77 | 0.78 |
| Salinity (surface) | Bottom depth | -0.75 | -0.77 | -0.77 | -0.78 | -0.79 | -0.79 | -0.75 | -0.77 | -0.78 | -0.77 | -0.79 | -0.78 |
| Salinity (depth) | Bottom depth | -0.76 | -0.77 | -0.76 | -0.77 | -0.76 | -0.76 | -0.74 | -0.72 | -0.72 | -0.74 | -0.75 | -0.76 |

Table SF6.2: Variables used in final models with VIF scores. Note January and February models were not run due to low number of presence points/sampling years.

| | **Jan** | **Feb** | **Mar** | **Apr** | **May** | **Jun** | **Jul** | **Aug** | **Sept** | **Oct** | **Nov** | **Dec** |
|---|---|---|---|---|---|---|---|---|---|---|---|---|
| **Temp (depth)** | 2.17 | 2.64 | 2.26 | 1.86 | 1.74 | 1.50 | 1.54 | 1.54 | 1.43 | 1.50 | 1.60 | 1.52 |
| **Salinity (depth)** | 2.55 | 2.22 | 1.98 | 1.56 | 1.38 | 1.27 | 1.33 | 1.32 | 1.19 | 1.27 | 1.49 | 1.87 |
| **O2 (depth)** | 2.46 | 2.38 | 2.27 | 2.09 | 1.79 | 1.54 | 1.52 | 1.32 | 1.22 | 1.54 | 1.73 | 2.18 |
| **Chl (surface)** | 1.74 | 1.57 | 1.30 | 1.09 | 1.38 | 1.42 | 1.21 | 1.14 | 1.21 | 1.42 | 1.31 | 1.44 |
| **NAO (sample month)** | 1.32 | 1.01 | 1.44 | 3.30 | 1.67 | 1.18 | 2.93 | 1.38 | 2.02 | 1.18 | 1.36 | 1.84 |
| **NAO (previous month)** | - | - | 1.43 | 1.69 | 4.31 | 1.39 | 2.50 | 1.71 | 1.75 | 1.39 | 1.21 | 1.37 |
| **NAO (previous winter)** | 1.32 | - | 1.52 | 4.10 | 3.69 | 1.06 | 1.64 | 1.34 | 1.64 | 1.06 | 1.07 | 1.53 |
| **AMO (sample month)** | - | - | 1.44 | 2.27 | 2.51 | 1.53 | 1.30 | 1.66 | 1.20 | 1.53 | 1.63 | 1.25 |
| **AMO (previous winter)** | - | - | 1.09 | 1.24 | 1.16 | 1.06 | 1.60 | 1.16 | 1.73 | 1.06 | 1.69 | 1.30 |


NOT PEER REVIEWED
