## Supplementary File 7 for "Modelling the spatial-temporal distributions and associated determining factors of a keystone pelagic fish"

### Supplementary File 7: Model performance metrics.

We used four metrics to quantify the performance of the monthly models (Table SF7). Testing and training area under the curve (AUC) scores are derived from receiver operator characteristic (ROC) analysis (Equation 1). The AUC is independent of threshold and prevalence (Phillips *et al.*, 2006), indicates the probability that a randomly chosen presence location ranks higher in suitability than a randomly chosen background point. Values range from 0 to 1. Values closer to 1 indicate the discriminatory power is good, whereas values close to and less than 0.5 indicate discriminatory power is poor (Fielding and Bell, 1997).

$$AUC = \frac{1}{n_A * n_N} \sum_{1}^{n_A} \sum_{1}^{n_N} S(TP, FP)$$

Equation 1: Both testing and training AUC scores are calculated in the same manor but test different subsections of the data (testing and training datasets respectively). $n_A$ = number of presences; $n_N$ = number of absences; $TP$ = number of true presences predicted by the model; $FP$ = number of false presences predicted by the model. The value of $S$ is conditional: If $TP > FP, S$ = 1; If $TP = FP, S$ = 0.5; If $TP < FP, S$ = 0. (Vida, 1993)

The True Skills Statistic (TSS) is a prevalence independent measure related to the widely used Cohens Kappa statistic which indicates the extent to which observed and predicted values are higher than expected by chance (Allouche *et al.*, 2006) (Equation 2). Values range from -1 to +1, with scores of +1 indicating perfect agreement and scores of 0 and less indicating values are no better than chance.

$$Sensitivity = \frac{TP}{TP + FA}$$

$$Specificity = \frac{TA}{FP + TA}$$

$$TSS = sensitivity + specificity - 1$$

Equation 2: TSS if a function of sensitivity (the probability of a model correctly predicting a presence) and specificity (the probability of a model correctly predicting an absence). $TP$ = the number presences accurately predicted; $FP$ = the number of presences falsely predicted (type 1 error); $TA$ = the number of absences accurately predicted; FA = the number of absences falsely predicted (type 2 error).

Finally, the Continuous Boyce Index (CBI) (Boyce *et al.*, 2002) (Equation 3) measures the degree to which model predictions differ from random over a moving window. It is independent of threshold and prevalence and is considered a robust measurement for presence-absence models. As with the TSS, CBI values range from -1 to +1, with scores of +1 indicating perfect agreement and scores of 0 and less indicating values are no better than chance and poor model performance.

NOT PEER REVIEWED

$$P_i = \frac{p_i}{\sum_{j=1}^{b} p_j}$$

$$E_i = \frac{a_i}{\sum_{j=1}^{b} a_j}$$

$$F_i = \frac{P_i}{E_i}$$

Equation 3: The Continuous Boyce Index. Distribution ranges are sub divided into $b$ bins. For each bin ($i$), both the predicted frequency of presence points ($P_i$) and the expected frequency of presence points if they were randomly distribution in the window ($E_i$) are calculated, and then used to create a predicted-to-expected ratio ($F_i$). $p_i$ = number of presence points predicted to fall into the distribution bin $i$; $\sum p_j$ = the total number of presence points; $a_i$ = the number of grid cells belonging to distribution class $i$; $\sum a_j$ = the total number of cells.

The AUC and TSS scores were obtained from the R package Dismo (Hijmans *et al.*, 2017) whilst the CBI was obtained with the R package enmSdm (Smith, 2019). For each month, each metric was calculated separately for each temporal fold and then averaged to produce average predictive performances.

Table SF7: Performance metrics for each of the monthly models. The mean represents the values obtained for all temporal fold, whereas the minimum and maximum values represent the highest and lowest scoring temporal fold. For all metrics, values range from -1 to +1, with scores of +1 indicating perfect agreement, and scores of -1 indicating completely imperfect agreement. Scores of 0 indicate values are no better than chance.

| | | **Mar** | **Apr** | **May** | **Jun** | **Jul** | **Aug** | **Sep** | **Oct** | **Nov** | **Dec** |
|---|---|---|---|---|---|---|---|---|---|---|---|
| **Train AUC** | **Mean** | 0.95 | 0.95 | 0.91 | 0.91 | 0.93 | 0.93 | 0.94 | 0.92 | 0.92 | 0.95 |
| | **Max** | 0.96 | 0.96 | 0.93 | 0.92 | 0.93 | 0.94 | 0.94 | 0.92 | 0.93 | 0.96 |
| | **Min** | 0.94 | 0.94 | 0.90 | 0.91 | 0.92 | 0.93 | 0.94 | 0.90 | 0.92 | 0.95 |
| **Test AUC** | **Mean** | 0.95 | 0.93 | 0.92 | 0.96 | 0.92 | 0.99 | 0.95 | 0.89 | 0.96 | 0.97 |
| | **Max** | 0.98 | 0.99 | 0.98 | 0.98 | 0.98 | 0.87 | 0.99 | 0.97 | 1.00 | 0.99 |
| | **Min** | 0.87 | 0.84 | 0.78 | 0.94 | 0.69 | 0.63 | 0.74 | 0.67 | 0.93 | 0.93 |
| **TSS** | **Mean** | 0.83 | 0.79 | 0.75 | 0.81 | 0.77 | 0.74 | 0.85 | 0.72 | 0.87 | 0.89 |
| | **Max** | 0.94 | 0.97 | 0.98 | 0.85 | 0.94 | 0.95 | 0.93 | 0.94 | 1.00 | 0.96 |
| | **Min** | 0.71 | 0.53 | 0.50 | 0.75 | 0.33 | 0.43 | 0.55 | 0.37 | 0.73 | 0.80 |
| **CBI** | **Mean** | 0.99 | 0.98 | 1.00 | 1.00 | 0.96 | 0.97 | 1.00 | 0.99 | 1.00 | 1.00 |
| | **Max** | 1.00 | 1.00 | 1.00 | 1.00 | 0.99 | 0.99 | 1.00 | 1.00 | 1.00 | 1.00 |
| | **Min** | 0.98 | 0.95 | 0.99 | 1.00 | 0.94 | 0.90 | 0.99 | 0.99 | 1.00 | 0.99 |

NOT PEER REVIEWED

NOT PEER REVIEWED
