## Supplementary File 8 for "Modelling the spatial-temporal distributions and associated determining factors of a keystone pelagic fish"

### Supplementary File 8:  Concatenating depth layers into single raster files for prediction.

Species distribution models including Maxent operate on a 2-dimensional surface. For each temporal fold used in the Maxent modelling, we concatenated each of the oceanographic variable depth layers into a single variable layer. These concatenated layers were used for predicting the estimated probability of presence of capelin (Bentlage *et al.*, 2013) (Figure SF8)

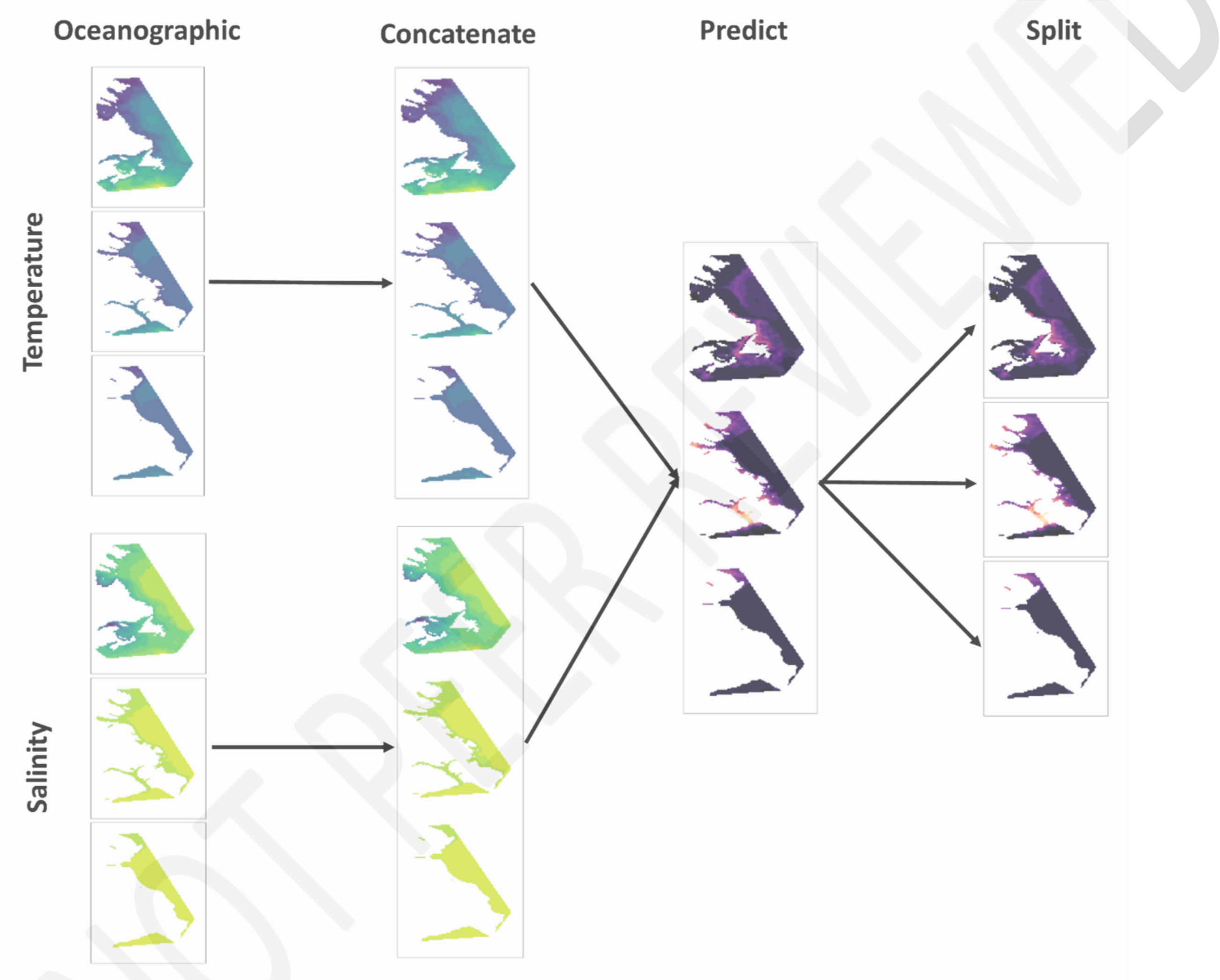


NOT PEER REVIEWED

Figure SF8: Example of transforming multiple 2-dimensonal oceanographic layers into a single 2-dimensional layer for prediction purposes. In this example, separate temperature depth layers for July 2004 are concatenated into a single ‘temperature layer’, and separate salinity depth layers for July 2004 concatenated into a single ‘salinity layer’. These concatenated layers are used to for prediction. The prediction layer is then split into depth layers. The year, month, depths, and oceanographic variables have been arbitrarily chosen for illustrative purposes only.
