## Supplementary File 9 for "Modelling the spatial-temporal distributions and associated determining factors of a keystone pelagic fish"

### Supplementary File 9: Year-month distribution of capelin presence points.

Table SF9: Number of presences available for modelling. Duplicates and points missing depth/environmental data have been removed, and presences have been reduced to one per environmental cell. Note January was not modelled due to the low number of years, and February was not modelled due to the low number of presence points.

| | Jan | Feb | Mar | Apr | May | Jun | Jul | Aug | Sep | Oct | Nov | Dec |
|---|---|---|---|---|---|---|---|---|---|---|---|---|
| **1998** | 0 | 0 | 0 | 36 | 53 | 103 | 20 | 0 | 28 | 53 | 120 | 8 |
| **1999** | 0 | 0 | 44 | 19 | 45 | 110 | 23 | 0 | 48 | 6 | 91 | 40 |
| **2000** | 0 | 0 | 40 | 24 | 45 | 96 | 19 | 0 | 51 | 24 | 112 | 54 |
| **2001** | 0 | 5 | 22 | 31 | 53 | 73 | 13 | 0 | 26 | 17 | 65 | 90 |
| **2002** | 0 | 0 | 32 | 42 | 59 | 89 | 17 | 0 | 68 | 28 | 48 | 59 |
| **2003** | 38 | 0 | 22 | 38 | 67 | 97 | 17 | 0 | 17 | 27 | 42 | 69 |
| **2004** | 73 | 0 | 0 | 29 | 58 | 96 | 25 | 24 | 39 | 15 | 75 | 71 |
| **2005** | 12 | 4 | 28 | 13 | 39 | 85 | 32 | 45 | 36 | 39 | 94 | 26 |
| **2006** | 31 | 1 | 46 | 11 | 2 | 103 | 22 | 47 | 33 | 48 | 78 | 47 |
| **2007** | 0 | 0 | 18 | 27 | 36 | 79 | 58 | 50 | 47 | 24 | 76 | 50 |
| **2008** | 0 | 0 | 26 | 18 | 54 | 105 | 12 | 54 | 39 | 24 | 92 | 39 |
| **2009** | 0 | 0 | 10 | 34 | 106 | 43 | 8 | 0 | 40 | 21 | 109 | 24 |
| **2010** | 0 | 0 | 15 | 18 | 48 | 102 | 25 | 7 | 57 | 48 | 101 | 27 |
| **2011** | 0 | 0 | 0 | 30 | 71 | 82 | 11 | 7 | 34 | 42 | 72 | 34 |
| **2012** | 0 | 0 | 0 | 0 | 0 | 0 | 6 | 9 | 39 | 0 | 0 | 0 |
| **2013** | 0 | 0 | 0 | 0 | 0 | 0 | 25 | 3 | 30 | 3 | 1 | 0 |
| **2014** | 0 | 0 | 0 | 0 | 0 | 0 | 0 | 6 | 61 | 3 | 0 | 0 |
| **Total** | 154 | 10 | 303 | 370 | 736 | 1263 | 333 | 252 | 693 | 422 | 1176 | 638 |

NOT PEER REVIEWED
