## Supplementary File 10 for "Modelling the spatial-temporal distributions and associated determining factors of a keystone pelagic fish"

### Supplementary File 10: Variation in Environmental Correlates.

The background points provide an overview of environmental conditions across the MCP area used in the models (Table SF10). Across all months, median chlorophyll concentration values ranged from 0.25 (mmol.m$^{-3}$) (August) to 1.1 (mmol.m$^{-3}$) (May) (smallest range in (1.63 mmol.m$^{-3}$), largest range in July (4.53 mmol.m$^{-3}$)). Median dissolved oxygen values ranged from 289.81 mmol.m$^{-3}$ (September) to 321.71 mmol.m$^{-3}$ (May) (smallest range in Oct (380.98 mmol.m$^{-3}$), largest range in July (446.14 mmol.m$^{-3}$)). The median salinity value was lowest in August (33.41 PSU) and highest in March (33.71 PSU) (smallest range in December (14.63 PSU), largest range in June (22.26 PSU)). Temperature ranged from a Median of 27.11 kelvin (March) to 277.67 kelvin (August) (smallest range in April (15.63 kelvin), largest range in August (25.88 kelvin)). Median AMO values during 'sampling month' ranged from 0.09 (April and May) to 0.14 (March, July, and August), whilst NAO values ranged from -0.75 (March) to 0.29 (May). The NAO median values from the previous month were lowest in July (-0.23) and highest in March (0.95). Median AMO winter values ranged from 0.09 (April and May) to 0.14 (March, July, and August) whilst NAO winter values ranged from -0.18 (July, August, September) to 0.37 (March). Table SF10 shows the maximum, minimum, range, mean, and median values of all environmental variables (oceanographic and climate oscillations) from the background points used in each of the monthly models.

Figure SF10 provides a visual representation of the range and distribution of the oceanographic variables used in the model (chlorophyll, dissolved oxygen, salinity, and temperature) across three months (March, June, and December) in the year 1998 at a single depth layer (11.8 meters).

Table SF10: Summary data of the environmental correlates associated with the background points used in each monthly model.

| | **Chlorophyll Concentration at surface (mmol.m$^{-3}$)** | | | | | **Dissolved Oxygen at sampling depth (mmol.m$^{-3}$)** | | | | |
|---|---|---|---|---|---|---|---|---|---|---|
| **Month** | **Min** | **Max** | **Mean** | **Median** | **Range** | **Min** | **Max** | **Mean** | **Median** | **Range** |
| **3** | 0.02 | 2.54 | 0.72 | 0.73 | 2.52 | 1.01 | 396.47 | 313.53 | 314.48 | 395.46 |
| **4** | 0.02 | 2.85 | 0.94 | 0.94 | 2.83 | 1.01 | 405.01 | 315.08 | 318.06 | 403.99 |
| **5** | 0.09 | 2.99 | 0.99 | 1.1 | 2.9 | 1.00 | 417.36 | 312.46 | 321.71 | 416.36 |
| **6** | 0.05 | 3.42 | 0.73 | 0.79 | 3.37 | 1.00 | 418.55 | 305.63 | 309.86 | 417.55 |
| **7** | 0.03 | 4.56 | 0.39 | 0.36 | 4.53 | 1.00 | 447.13 | 297.81 | 297.8 | 446.14 |
| **8** | 0.03 | 1.68 | 0.26 | 0.25 | 1.65 | 0.99 | 402.95 | 290.45 | 290.48 | 401.96 |
| **9** | 0.03 | 2.69 | 0.29 | 0.26 | 2.66 | 0.99 | 394.89 | 287.48 | 289.81 | 393.9 |
| **10** | 0.03 | 2.37 | 0.37 | 0.35 | 2.34 | 0.94 | 381.93 | 289.12 | 292.27 | 380.98 |
| **11** | 0.04 | 2.52 | 0.4 | 0.36 | 2.48 | 0.96 | 386.5 | 295.34 | 300.9 | 385.53 |
| **12** | 0.09 | 2.22 | 0.39 | 0.37 | 2.13 | 0.95 | 390.24 | 301.35 | 306.65 | 389.29 |
| | **Salinity Concentration at sampling depth (PSU)** | | | | | **Temperature at sampling depth (kelvin)** | | | | |
| **Month** | **Min** | **Max** | **Mean** | **Median** | **Range** | **Min** | **Max** | **Mean** | **Median** | **Range** |
| **3** | 20.25 | 35.69 | 33.33 | 33.71 | 15.44 | 270.68 | 286.76 | 275.05 | 275.11 | 16.07 |
| **4** | 15.92 | 35.69 | 33.3 | 33.69 | 19.77 | 270.99 | 286.62 | 275.25 | 275.39 | 15.63 |
| **5** | 14.15 | 35.76 | 33.23 | 33.55 | 21.61 | 270.82 | 287.69 | 275.91 | 276.32 | 16.87 |
| **6** | 13.49 | 35.75 | 33.11 | 33.54 | 22.26 | 270.73 | 291.46 | 277.06 | 276.86 | 20.73 |
| **7** | 14.46 | 35.95 | 33.04 | 33.51 | 21.49 | 270.38 | 293.83 | 278.32 | 277.21 | 23.45 |

NOT PEER REVIEWED

| | | | | | | | | | | |
|---|---|---|---|---|---|---|---|---|---|---|
| **8** | 16.69 | 35.84 | 32.95 | 33.41 | 19.15 | 270.77 | 296.66 | 279.52 | 277.67 | 25.88 |
| **9** | 16.57 | 36 | 33.01 | 33.43 | 19.44 | 270.67 | 296.14 | 279.3 | 277.65 | 25.47 |
| **10** | 17.97 | 35.92 | 33.02 | 33.48 | 17.95 | 270.72 | 295.86 | 278.57 | 277.54 | 25.14 |
| **11** | 19.38 | 35.93 | 33.06 | 33.48 | 16.55 | 270.69 | 290.06 | 277.46 | 277.18 | 19.37 |
| **12** | 21.2 | 35.83 | 33.17 | 33.58 | 14.63 | 270.54 | 289.79 | 276.48 | 276.61 | 19.25 |
| | **AMO from previous winter** | | | | | **AMO from sampling month** | | | | |
| **Month** | **Min** | **Max** | **Mean** | **Median** | **Range** | **Min** | **Max** | **Mean** | **Median** | **Range** |
| **3** | -0.09 | 0.21 | 0.11 | 0.14 | 0.31 | -0.16 | 0.29 | 0.11 | 0.1 | 0.45 |
| **4** | -0.09 | 0.21 | 0.06 | 0.09 | 0.31 | -0.13 | 0.43 | 0.1 | 0.08 | 0.56 |
| **5** | -0.09 | 0.21 | 0.07 | 0.09 | 0.31 | -0.06 | 0.46 | 0.13 | 0.15 | 0.52 |
| **6** | -0.09 | 0.21 | 0.08 | 0.13 | 0.31 | -0.12 | 0.49 | 0.21 | 0.2 | 0.61 |
| **7** | -0.09 | 0.21 | 0.09 | 0.14 | 0.31 | -0.07 | 0.49 | 0.24 | 0.21 | 0.55 |
| **8** | -0.07 | 0.21 | 0.11 | 0.14 | 0.28 | 0.06 | 0.53 | 0.28 | 0.32 | 0.47 |
| **9** | -0.09 | 0.21 | 0.08 | 0.13 | 0.31 | 0.06 | 0.45 | 0.25 | 0.24 | 0.4 |
| **10** | -0.09 | 0.21 | 0.06 | 0.1 | 0.31 | -0.04 | 0.43 | 0.23 | 0.24 | 0.47 |
| **11** | -0.09 | 0.21 | 0.08 | 0.1 | 0.31 | -0.07 | 0.32 | 0.12 | 0.14 | 0.39 |
| **12** | -0.09 | 0.21 | 0.07 | 0.1 | 0.31 | -0.13 | 0.28 | 0.11 | 0.17 | 0.41 |
| | **NAO from previous winter** | | | | | **NAO from sampling month** | | | | |
| **Month** | **Min** | **Max** | **Mean** | **Median** | **Range** | **Min** | **Max** | **Mean** | **Median** | **Range** |
| **3** | -0.69 | 1.66 | 0.35 | 0.37 | 2.35 | -2.32 | 1.78 | -0.46 | -0.75 | 4.10 |
| **4** | -3.03 | 1.66 | 0.06 | 0.12 | 4.69 | -1.29 | 2.31 | 0.13 | 0.27 | 3.60 |
| **5** | -3.03 | 1.66 | 0.04 | 0.12 | 4.69 | -1.33 | 1.49 | 0.30 | 0.29 | 2.82 |
| **6** | -3.03 | 1.66 | 0.06 | 0.03 | 4.69 | -1.25 | 1.03 | -0.15 | -0.23 | 2.28 |
| **7** | -3.03 | 1.66 | 0.08 | -0.18 | 4.69 | -1.05 | 0.46 | -0.07 | 0.02 | 1.51 |
| **8** | -3.03 | 1.47 | -0.08 | -0.18 | 4.50 | -0.97 | 0.79 | -0.20 | -0.09 | 1.76 |
| **9** | -3.03 | 1.66 | -0.06 | -0.18 | 4.69 | -1.89 | 1.00 | -0.02 | 0.10 | 2.89 |
| **10** | -3.03 | 1.66 | -0.07 | 0.12 | 4.69 | -1.89 | 1.20 | -0.33 | -0.58 | 3.09 |
| **11** | -3.03 | 1.66 | 0.12 | 0.12 | 4.69 | -1.64 | 1.57 | 0.01 | 0.16 | 3.21 |
| **12** | -3.03 | 1.66 | 0.22 | 0.12 | 4.69 | -3.63 | 2.60 | -0.14 | -0.14 | 6.23 |

| | **NAO from month previous to sampling** | | | | |
|---|---|---|---|---|---|
| **Month** | **Min** | **Max** | **Mean** | **Median** | **Range** |
| **3** | -3.98 | 2.86 | 0.28 | 0.95 | 6.84 |
| **4** | -2.32 | 1.78 | 0.18 | 0.58 | 4.10 |
| **5** | -1.29 | 2.31 | 0.09 | 0.27 | 3.60 |
| **6** | -1.33 | 1.49 | 0.13 | 0.27 | 2.82 |
| **7** | -1.25 | 1.03 | -0.22 | -0.23 | 2.28 |
| **8** | -1.24 | 0.40 | -0.10 | -0.12 | 1.64 |
| **9** | -0.97 | 0.79 | -0.17 | -0.21 | 1.76 |
| **10** | -1.89 | 1.00 | -0.22 | 0.05 | 2.89 |
| **11** | -1.89 | 1.2 | -0.21 | -0.08 | 3.09 |
| **12** | -1.64 | 1.21 | 0.16 | 0.28 | 2.85 |

NOT PEER REVIEWED

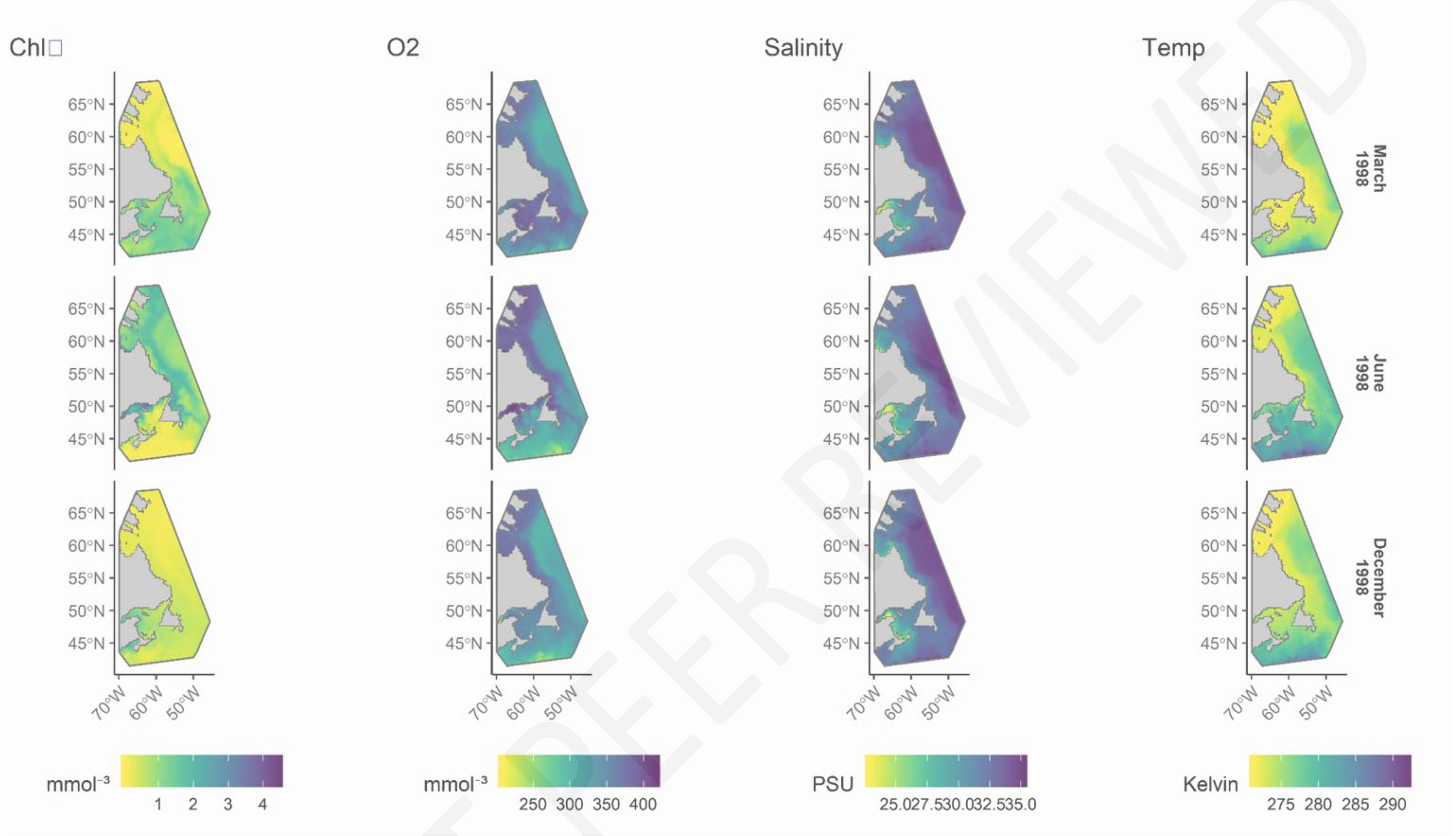


Figure SF10: Modelled oceanographic conditions at 11.8 meters during March, June, and December 1998. The year, month, and depths have been arbitrarily chosen for illustrative purposes only.

NOT PEER REVIEWED
